# Totally tubular: ASO-mediated knock-down of *G2019S*-Lrrk2 modulates lysosomal tubule-associated antigen presentation in macrophages

**DOI:** 10.1101/2023.07.14.549028

**Authors:** Rebecca L. Wallings, Julian R. Mark, Hannah A. Staley, Drew A. Gillett, Noelle Neighbarger, Holly Kordasiewicz, Warren D. Hirst, Malú Gámez Tansey

## Abstract

Genetic variation around the *LRRK2* gene affects risk of both familial and sporadic Parkinson’s disease (PD). LRRK2 levels have become an appealing target for potential PD-therapeutics with LRRK2 antisense oligonucleotides (ASOs) now in clinical trials. However, LRRK2 has been suggested to play a fundamental role in peripheral immunity, and it is currently unknown if targeting increased LRRK2 levels in peripheral immune cells will be beneficial or deleterious. Furthermore, the precise role of LRRK2 in immune cells is currently unknown, although it has been suggested that LRRK2-mediated lysosomal function may be crucial to immune responses. Here, it was observed that *G2019S* macrophages exhibited increased stimulation-dependent lysosomal tubule formation (LTF) and MHC-II trafficking from the perinuclear lysosome to the plasma membrane in an mTOR dependent manner with concomitant increases in pro-inflammatory cytokine release. Both ASO-mediated knock down of mutant *Lrrk*2 and LRRK2 kinase inhibition ameliorated this phenotype and decreased these immune responses in control cells. Given the critical role of antigen presentation, lysosomal function, and cytokine release in macrophages, it is likely LRRK2-targetting therapies may have therapeutic value with regards to mutant *LRRK2* but deleterious effects on the peripheral immune system, such as altered pathogen control and infection resolution.

## Introduction

Parkinson’s disease (PD) is a common progressive neurodegenerative disease, affecting around 1-2% of the population over the age of 65 [1]. Prevalence of PD is expected to increase two-fold by the year 2030 [2]. In addition, it is estimated that projected total economic burden will surpass $79 billion by 2037 [3], highlighting the need for interventions that could delay disease progression. The fact that there are currently no disease-modifying drugs for people with PD indicates that knowledge gaps still need to be closed to identify ways to cure or prevent this disease.

The most prevalent LRRK2 mutation, *G2019S*, resides in the kinase domain and causes a 2- 3-fold increase in kinase activity [4, 5]. Furthermore, increased LRRK2 mRNA and protein levels have been observed in B cells, T cells, non-classical CD16+ monocytes [6] and neutrophils [7] of patients with sporadic PD when compared to age-matched healthy controls. LRRK2 levels have therefore become an appealing target for potential PD-therapeutics, with *LRRK2* antisense oligonucleotides (ASOs) now in clinical trials. Indeed, in a preclinical mouse model, administration of *Lrrk2* ASOs to the brain reduces LRRK2 protein levels and fibril-induced α-synuclein inclusions [8]. Similarly, the use of an ASO that blocks splicing of *LRRK2*exon 41, which encodes part of the kinase domain, reverses aberrant endoplasmic reticulum (ER) calcium levels and mitophagy defects in PD patient-derived cell lines harbouring the *LRRK2 G2019S* mutation [9, 10]. The administration of *Lrrk2* ASOs have also been shown to rescue aberrant Rab10 phosphorylation levels and autophagic processing in the brains of mice expressing human *G2019S-LRRK2*[11]. Although these early studies show promising results with LRRK2 ASO in the brain, relatively little is known about the potential effects of targeting LRRK2 levels in the periphery, where it is highly expressed in immune cells [6, 12].

It has been shown that LRRK2 levels increase in immune cells upon immune cell activation [6, 13]. However, whether LRRK2 expression increases in peripheral immune cells to dampen or promote inflammation is still unknown. Of note, complete abolition of LRRK2 kinase activity in the peripheral immune system leads to deleterious effects in Lrrk2-KO models, with increased risk of infection and decreased pathogen control [12, 14–16]. This immune dysfunction may be mediated by lysosomal defects, as it has recently been demonstrated that a mouse *Lrrk2*-knockout macrophage cell line displays vacuolization and lipofuscin autofluorescence upon lysosomal overload stress [17]. Based on this, it can be inferred that LRRK2 may modulate lysosomal function in peripheral immune cells and increases in expression to regulate inflammation.

Upon toll-like receptor 4 (TLR4) stimulation with lipopolysaccharide (LPS), LRRK2 levels increase and the protein is recruited to endolysosomal membranes whereby it regulates the autophagy pathway in RAW264.7 macrophage cells and HEK-293 cells [18, 19]. Such data suggests that LRRK2 may increase in response to inflammatory stimuli to mediate lysosomal function. Interestingly, LRRK2 has recently been shown to mediate tubulation and vesicle sorting from lysosomes in astrocytes [20]. Bonet-Ponce and colleagues demonstrated that, upon lysosomal membrane-rupturing, LRRK2 is recruited to the lysosomal membrane whereby it mediates the formation of lysosomal tubules and release of lysosomal contents, with *G2019S*-Lrrk2 expression significantly increasing the formation of these tubules in astrocytes. Interestingly, it is known that lysosomal tubulation is usually observed in macrophages and other professional phagocytes undergoing immune activation [21]. Lysosomal tubules are crucial for two immune-related functions upon immune activation; phagocytosis and antigen presentation [21, 22]. This is intriguing as LRRK2 has been heavily implicated in modulating phagocytosis [23] and LRRK2 expression is also positively correlated with HLA-DR expression in human monocytes [6, 13], suggesting a potential role of LRRK2 in antigen presentation. It is therefore possible that LRRK2 mediates inflammatory responses, such as antigen presentation and phagocytosis, via lysosomal tubulation.

The aim of this study is two-fold; to assess the effects of Lrrk2 knock-down via ASOs on immune cell responses and lysosomal function, as well as to test the hypothesis that LRRK2 mediates inflammatory responses, specifically antigen presentation, via the formation of lysosomal tubules. Here, we employ *Lrrk2* ASOs and Lrrk2 kinase inhibitors to investigate a novel mechanism that may link the role of LRRK2 at the lysosome to its role in inflammation and antigen presentation. As both strategies are being evaluated in clinical trials with PD patients [8, 24], our inclusion of Lrrk2 knock-down and targeting of its kinase activity provides vital information on the potentially deleterious effects of reduced LRRK2 activity in peripheral immune cells. Here, we observe an increase in stimulation-dependent antigen presentation, and cytokine release in peritoneal macrophages (pMacs) from *G2019S* BAC mice relative to wild-type Lrrk2 over-expressing (WTOE) and B6 controls. We also observe alterations in lysosomal function, with increased pan-cathepsin activity and degradative capacity of lysosomes in *G2019S* pMacs early in the inflammatory response. Knock-down of *Lrrk2*, as well as Lrrk2 kinase-inhibition, successfully ameliorate these phenotypes, and nanostring-based transcriptomic profiling suggest altered vesicular trafficking, lysosomal positioning, and autophagy activity may underlie the effects of *Lrrk2* knock-down on antigen presentation. Indeed, it was observed that *G2019S* pMacs exhibited increased stimulation-dependent lysosomal tubule formation (LTF) and MHC-II trafficking from the perinuclear lysosome to the plasma membrane in an mTOR-dependent manner.

## Results

### *G2019S* BAC transgenic pMacs exhibit increased antigen presentation and cytokine release

LRRK2 has previously been shown to increase in immune cells from both murine pre-clinical models and patient cells in response to inflammatory stimuli[6, 13, 25–27]. To determine if Lrrk2 increases in response to such stimulus in pMacs, Lrrk2 protein levels were assessed in pMacs treated with 100U of IFNγ for 18h. An increase in Lrrk2 levels was observed in all three genotypes upon IFNγ treatment (Fig.1A, B). Furthermore, significantly more Lrrk2 is present in *WTOE* and *G2019S* pMacs relative to B6 pMacs in both vehicle- and IFNγ-treated pMacs, with no significant difference seen between the two BAC models. Furthermore, we observed that IFNγ-treatment caused a significant increase in phosphorylated Lrrk2 in all genotypes, indicative of increased kinase activity levels [28], with significantly higher levels of phosphorylated Lrrk2 observed in *G2019S*pMacs relative to the other genotypes (Sup.Fig1A, B). Co-treatment with 100nM of the LRRK2 kinase inhibitor, PF-06685360 (PF360), significantly reduced phosphorylated Lrrk2 levels in all genotypes and prevented stimulation-dependent increase (Sup. Fig1B).

**Figure 1.**
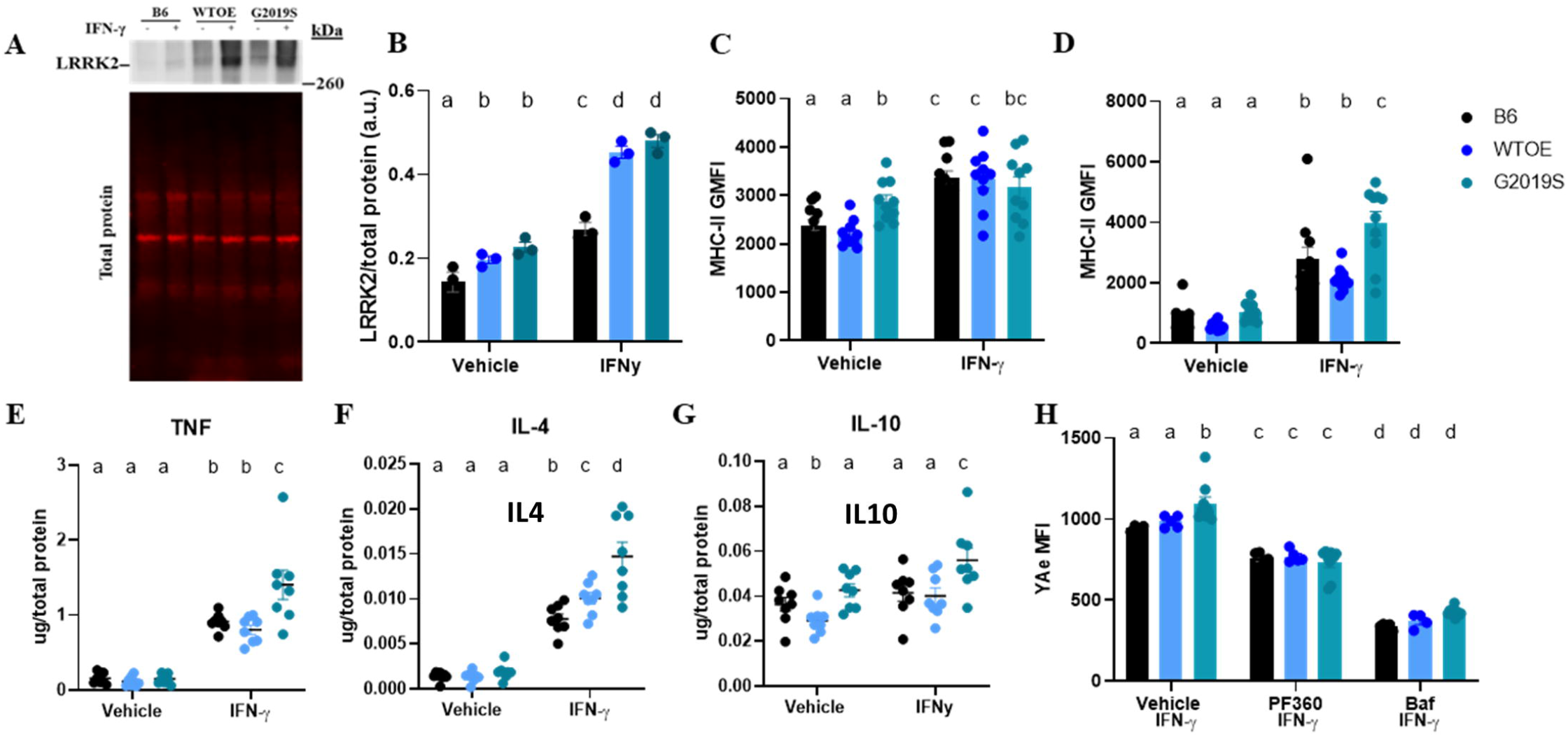
*G2019S*BAC transgenic pMacs exhibit increased antigen presentation and cytokine release: pMacs from 10-12-week-old male B6, *WTOE* or *G2019S* mice were stimulated with 100U IFNγ for 18-hours. (**A, B**) Lrrk2 protein levels were assessed and normalized to total protein levels and quantified. Representative western blots shown. (**C, D**) MHC-II GMFI was quantified on LPMs and SPMs via flow cytometry. (**E, F, G**) Levels of the cytokines TNF, IL4 and IL10 in media were assessed, normalized to total protein levels and quantified. (**H**) Cells were co-treated with 100nM PF360 or 50nM Bafilomycin A1 (baf) and Yae MFI was quantified on LPMs via flow-cytometry. Bars represent mean +/- SEM (N = 6-10). One/Two-way ANOVA, Bonferroni post-hoc, groups sharing the same letters are not significantly different (p>0.05) whilst groups displaying the same letter are significantly different (p<0.05).

pMacs are not a homogenous population of cells, but rather a mix of small and large pMacs (SPMs and LPMs, respectively). LPMs from *G2019S* mice, which are distinguished from SPMs based on CD11b expression (Sup. Fig. 1C) [29] are resident to the peritoneal cavity and are traditionally thought of as anti-inflammatory and phagocytic [29]. SPMs, on the other hand, are generated from bone-marrow-derived myeloid precursors which migrate to the peritoneal cavity in response to infection, inflammatory stimuli, or thioglycolate, and present a pro-inflammatory functional profile [29]. When assessing LPM and SPM count, it was observed that no differences were seen in LPM count between genotype nor treatment (Sup.Fig.1D), although a significant upregulation of SPMs was observed in *WTOE* pMacs upon IFNγ treatment relative to G2019S and B6 pMacs (Sup.Fig.1E). As bone-marrow-derived myeloid precursors are known to differentiate into SPMs upon inflammation, it may be, therefore, *WTOE* bone-marrow-derived myeloid precursors present in these cultures have increased propensities to differentiate in such conditions.

To begin to investigate the effects of *Lrrk2-* over-expression and mutation on pMac function, we assessed MHC-II expression on both LPMs and SPMs as a measure of antigen presentation. Although no significant differences were observed between genotypes regarding MHC-II+ LPM count (Sup.Fig.1F), it was observed that *G2019S-* expressing LPMs express significantly more MHC-II in vehicle-treated LPMs, with all three genotypes increasing MHC-II expression with IFNγ treatment (Fig.1C). In SPMs, no differences between genotypes were observed in vehicle treatment, however increased MHC-II expression was observed in IFNγ-treated *G2019S*-expressing SPMs relative to the other genotypes (Fig.1D). Regarding MHC-II+ SPM count, increased counts were observed in IFNγ-treated WTOE pMacs (Sup.Fig.1G) however when calculated as a percentage of total SPMs, this difference did not persist (Sup.Fig.1H) suggesting this observation was due to more SPMs overall as opposed to more MHC-II+ SPMs.

To see if alterations in MHC-II expression were accompanied by changes in cytokine release, media from vehicle-and IFNγ treated cells were collected and cytokine levels quantified. Increased levels of the pro-inflammatory TNF (Fig. 1E) and the anti-inflammatory IL4 were observed in media from IFNγ-treated *G2019S* pMacs (Fig.1F). As well, increased levels of the anti-inflammatory IL10 were also observed in media from *G2019S* pMacs when treated with vehicle relative to media from WTOE and B6 pMacs. In both *WTOE* and *G2019S* pMacs, IFNγ treatment caused a reduction in IL10 in the media compared to vehicle treatment, however no change was observed in B6 pMacs (Fig. 1G). No significant differences were observed in other cytokines measured (Sup.Fig.1I-N).

The Eα: YAe model, where presentation of the Eα peptide on MHC-II may be detected by flow cytometry using the YAe antibody (Sup. Fig.2), allows us to measure antigen presentation of a peptide directly and acts as a measure of the whole antigen presentation pathway, from uptake to peptide loading to presentation. It was observed here that Yae MFI was significantly upregulated in IFNγ-treated *G2019S*-expressing LPMs relative to the other genotypes (Fig.1H). When co-treated with PF360, this phenotype was ameliorated and no significant differences between genotypes observed. Antigen presentation and pathogen sensing requires protease action and sufficient lysosomal function in order to occur [30], which is why lysosomotropic agents have been shown to decrease peptide-loaded-MHC-II surface expression in antigen presenting cells [31]. Indeed, when co-treated with the vacuolar H+ ATPase (V-ATPase) inhibitor, Bafilomycin A1, Yae MFI significantly decreased in all three genotypes. Due to this observation and the crucial role of the lysosome in antigen presentation, we next sought to probe the effects of Lrrk2 over-expression and mutation on lysosomal function in these pMacs.

### LRRk2 kinase activity modulates lysosomal function early in the inflammatory response in cells engaging in antigen presentation

The immune response to an inflammatory stimulus is a dynamic process with peaks in different cellular activities occurring at different times. Although LRRK2 levels have been shown to increase in response to inflammatory stimuli, reports typically measure LRRK2 levels at end time-point, neglecting to show changes in LRRK2 levels over time during the inflammatory response. We therefore wanted to examine how LRRK2 levels and phosphorylation change over time in response to IFNγ. Regarding total Lrrk2 levels, a significant, steady increase in LRRK2 levels were seen over the 18-hour IFNγ treatment with *G2019S* and *WTOE*-expressing pMacs consistently expressing increased levels relative to B6 pMacs (Fig 2A, B). Regarding phosphorylated Lrrk2, a similar pattern was observed, with levels of pLRRK2 at S935 increasing over the 18-hour IFNγ treatment in all genotypes, with increased levels observed in *G2019S* pMacs relative to the other genotypes at the 18-hour time point (Fig.2C).

**Figure 2.**
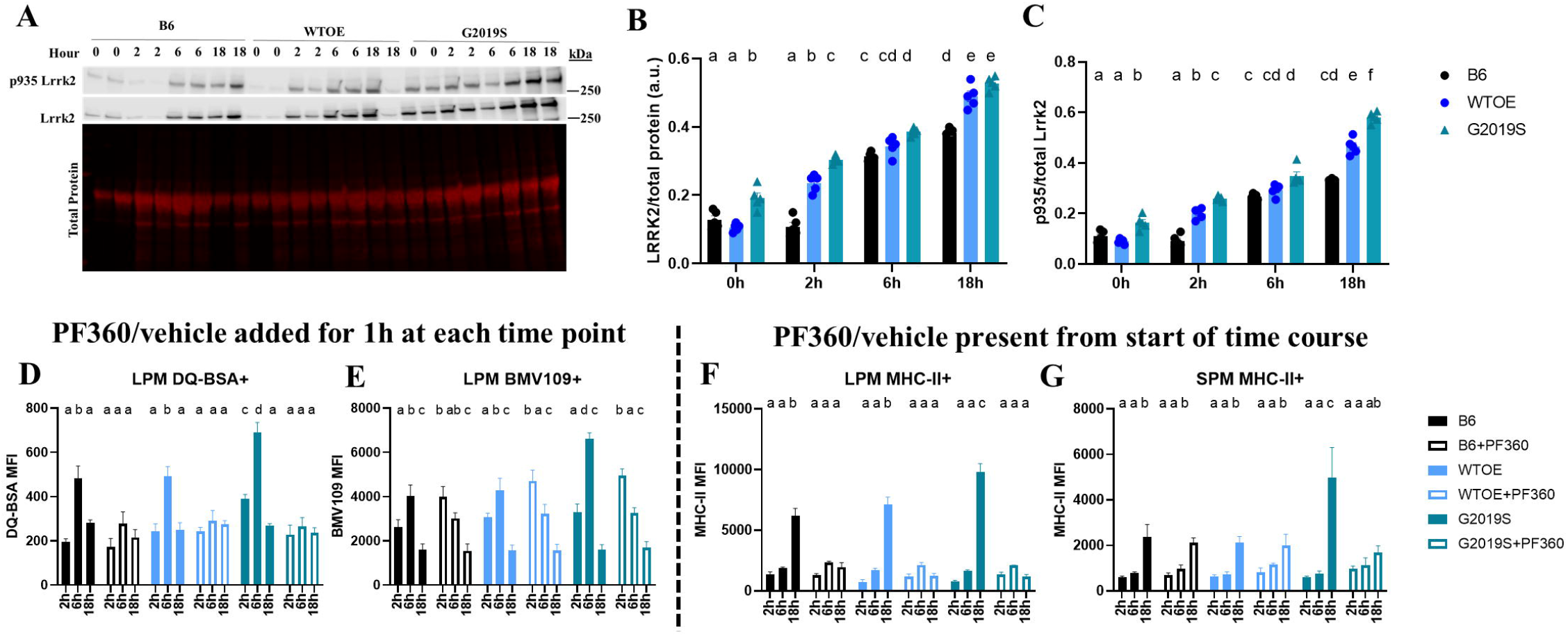
LRRk2 kinase activity modulates lysosomal function early in the inflammatory response in cells engaging in antigen presentation: pMacs from 10- 12-week-old male B6, *WTOE*or *G2019S* mice were stimulated with 100U IFNγ and harvested at 2-, 6- or 18-hours. (**A, B, C**) Lrrk2 protein and phosphorylated protein levels were assessed and normalized to total protein/total Lrrk2 levels and quantified. Representative western blots shown. pMacs from 10-12-week-old male B6, *WTOE* or *G2019S* mice were stimulated with 100U IFNγ and harvested at 2-, 6- or 18-hours after 1-hour treatment with 100nM PF360 immediately prior to harvesting. (**D**) DQ-BSA MFI was quantified in LPMs via flow cytometry. (**E**) BMV109 MFI was quantified in LPMs via flow cytometry. pMacs from 10-12-week-old male B6, *WTOE* or *G2019S* mice were stimulated with 100U IFNγ +/- PF360 and harvested at 2-, 6- or 18-hours. (**F, G**) MHC-II GMFI was quantified in SPMs and LPMs via flow cytometry. Bars represent mean +/- SEM (N = 8-10). Three-way ANOVA, Bonferroni post-hoc, groups sharing the same letters are not significantly different (p>0.05) whilst groups displaying the same letter are significantly different (p<0.05).

Next, pMacs were plated in the presence of IFNγ and collected at 2-, 6- or 18-hours into the treatment and measures of lysosomal activity were recorded via flow cytometry. To understand the role of Lrrk2-kinase activity over this time course, we applied one of two experimental designs (Sup. Fig. 3A); PF360 or vehicle was either applied for 1-hour prior to collection at each time point, or, PF360 or vehicle was present from the start of the IFNγ treatment when cells were plated. In the conditions where PF360 or vehicle was present for the 1-hour prior to collection, it was observed that lysosomal degradation, as measured by DQ Red BSA fluorescence, peaked at 6-hour into the IFNγ-treated LPMs in vehicle conditions for all 3 genotypes (Fig. 2D). *G2019S*-pMacs treated with vehicle exhibited even greater lysosomal degradation at this 6-hour time point than WTOE and B6 pMacs. Interestingly, the 1-hour treatment with PF360 prior to collection significantly reduced this 6-hour peak in lysosomal degradation in all 3 genotypes. Similarly, cathepsin activity was measured using the pan-cathepsin probe, BMV109, and a significant peak in cathepsin activity was observed at the 6-hour time point in vehicle-treated pMacs, independent of genotype, relative to the other time points (Fig. 2E). Interestingly, *G2019S*-expressing LPMs had significantly increased cathepsin activity at this 6-hour time point relative to the other genotypes. Treatment of PF360 for 1-hour prior to collection could decrease this 6-hour peak in all genotypes, mirroring what was seen in the DQ Red BSA measurements. However, it was also observed that this PF360 treatment significantly increased cathepsin activity at the 2-hour time point in all 3 genotypes. Interestingly, when measuring MHC-II expression on LPMs and SPMs, although effects of genotype were observed at the 18-hour time point, no significant effects of 1-hour PF360 treatment were observed at any time point (Sup.Fig.3B, C).

Keeping in mind that the role of the lysosome occurs earlier in the inflammatory response than the final end-product of antigen presentation [30], we repeated these experiments with PF360 present from the start of the 18-hour IFNγ treatment. Interestingly, in this condition, the increased MHC-II expression in *G2019S* LPMs and SPMs at 18-hours was ameliorated (Fig. 2F, G). MHC-II expression was also significantly decreased at the 18-hour time point in B6 and *WTOE* LPMs treated with PF360 from the start of the IFNγ response (Fig. 2G). Similar changes in DQ Red BSA and BMV109 fluorescence were observed in PF360-treated pMacs in this experimental design that were seen in the 1-hour PF360 condition (Sup. Fig. 3D, E). Collectively, such data suggest that Lrrk2 kinase activity modulates antigen presentation earlier in the inflammatory response, as PF360 was unable to ameliorate the *G2019S*-mediated increase in MHC-II on LPMs when present for 1-hour prior to collection at the 18-hour time point, but could decrease MHC-II expression when present from the start of the inflammatory response. Given what is understood about the role of the lysosome in antigen presentation pathways, and the fact that we see significant effects of Lrrk2-kinase inhibition on lysosomal activity at the 6-hour time point, we hypothesized that Lrrk2 may regulate antigen presentation via lysosomal activity early in the inflammatory response (Sup.Fig.3F).

### *G2019S* pMac antigen presentation and lysosomal phenotypes are rescued by knock-down of *Lrrk2*

It is currently unknown whether targeting increased LRRK2 levels in peripheral immune cells with Lrrk2-targetting ASOs will be beneficial or deleterious to immune cell function, therefore these flow cytometry-based assays were repeated in pMacs nucleofected with *Lrrk2*-targettng ASO (Ionis) or scramble control. Ionis provided 3 *Lrrk2*-targetting ASOs which were used to nucleofect B6 pMacs at a concentration of 1µg per reaction (2 × 10^6^ cells per reaction); it was observed that ASO 3 significantly reduced total Lrrk2 levels in pMacs relative to scramble control at both the protein and the mRNA level (Sup.Fig.4A, B). When pMacs from BAC mice and B6 controls were nucleofected with this *Lrrk2* ASO, a significant reduction in Lrrk2 protein levels were observed in all genotypes relative to scramble ASO controls (Fig.3A, B). Furthermore, in *Lrrk2* ASO-nucleofected cells, 18-hour treatment with IFNγ was unable to increase Lrrk2 levels as seen in scramble controls.

**Figure 3.**
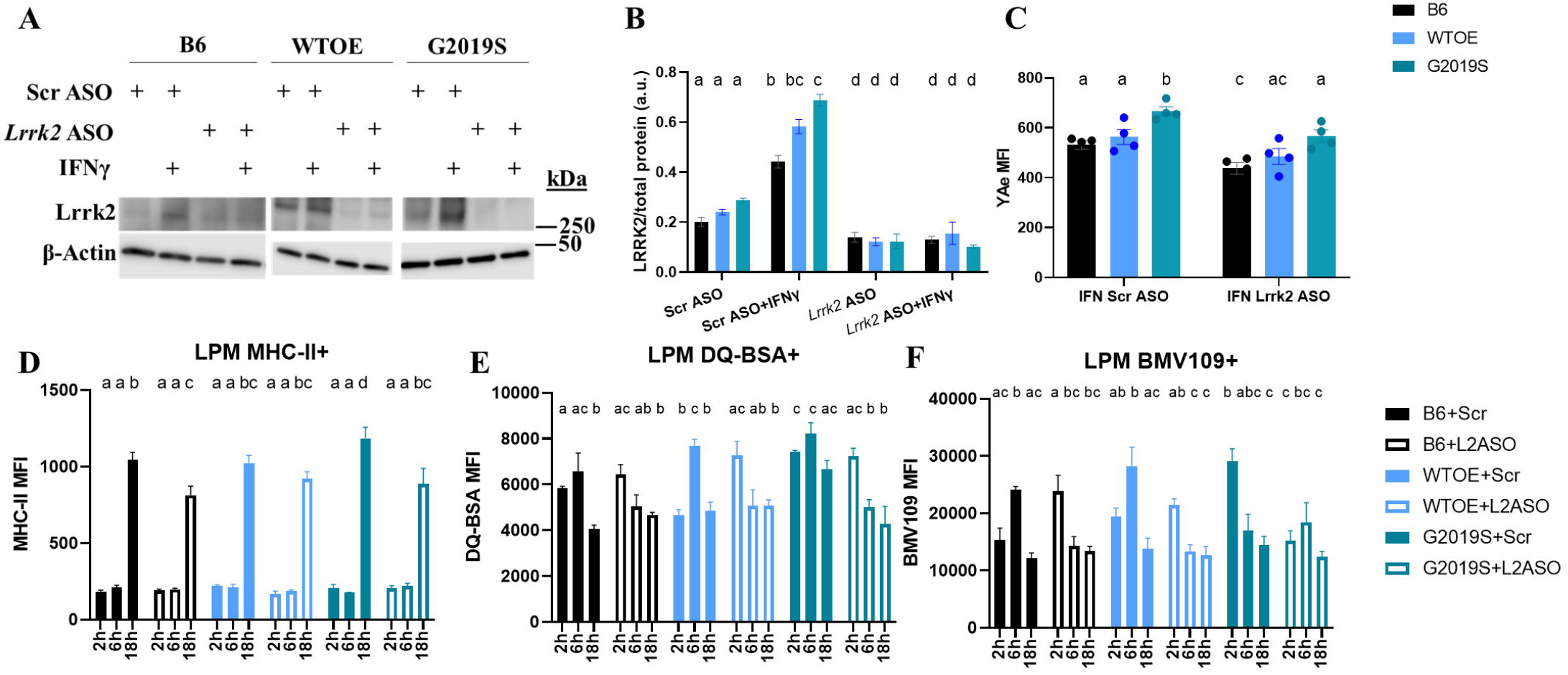
*G2019S* pMac antigen presentation and lysosomal phenotypes are rescued by knock-down of LRRK2: pMacs from 10-12-week-old male B6, *WTOE* or *G2019S* mice were nuclefoected with a *Lrrk2*-targetting ASO or scramble control and stimulated with 100U IFNγ and harvested at 2-, 6- or 18-hours. (**A, B**) Lrrk2 protein levels normalized to β-actin levels and quantified. (**C**) YAe MFI was quantified on LPMs via flow-cytometry. (**D**) MHC-II GMFI was quantified in LPMs via flow cytometry. (**E**) DQ-BSA MFI was quantified in LPMs via flow cytometry. (**F**) BMV109 MFI was quantified in LPMs via flow cytometry. Bars represent mean +/- SEM (N = 8-10). Two/Three-way ANOVA, Bonferroni post-hoc, groups sharing the same letters are not significantly different (p>0.05) whilst groups displaying the same letter are significantly different (p<0.05).

Using the Eα: YAe model, antigen presentation was assessed in nucleofected pMacs after 18- hours of IFNγ treatment. It was seen that in scramble conditions increased YAe MFI was observed in *G2019S*-expressing LPMs relative to other genotypes as previously shown (Fig.3C). Interestingly, nucleofection with the *Lrrk2-* targetting ASO significantly reduced this YAe MFI in *G2019S*-expressing LPMs. Furthermore, knock-down of *Lrrk2* also significantly decreased YAe MFI in B6 pMacs, with a trend seen in WTOE pMacs. This was supported by flow cytometry-based MHC-II expression analysis in LPMs (Fig. 3D).

In order to assess the effects of Lrrk2 knock-down on lysosomal function, lysosomal degradation and cathepsin activity was measured via flow cytometry as before. Similar to what was observed in experiments assessing effects of Lrrk2-kinase inhibition, *Lrrk2* ASO treatment caused a significant reduction in both lysosomal protein degradation (Fig.3E) and cathepsin activity (Fig.3F) at the 6-hour time-point in *G2019S* and *WTOE* LPMs. All genotypes had a peak in lysosomal degradation at 6-hour time point when treated with scramble ASO, however interestingly treatment with *Lrrk2* ASO caused the peak to appear earlier at the 2-hour time point instead (Fig.3E).

### Lrrk2 kinase inhibition and knock-down via ASO reduces trafficking of MHC-II to the plasma membrane

As knock-down of Lrrk2 and Lrrk2-kinase inhibition reduces both YAe MFI and MHC-II levels via flow cytometry suggests there is decreased transport of peptide-loaded MHC-II-complexes from the lysosome to the cell surface. It can therefore be hypothesized that an increase in intracellular MHC-II expression would be observed in conditions where Lrrk2 is knocked-down, or its kinase activity inhibited. To test this, using pMacs treated with the LRRK2-kinase inhibitor, PF360, or nucleofected with a scramble or *Lrrk2*-ASO, we co-stained for extracellular vs intracellular MHC-II (exMHCII vs icMHCII) and monitored expression via fluorescent microscopy. It was observed that *G2019S* pMacs exhibited increased Ex:IcMHCII ratio relative to other genotypes, which was ameliorated upon LRRK2 kinase inhibition (Fig.4A, B), with this treatment decreasing Ex:IcMHCII ratio in all genotypes. Such data suggests that Lrrk2-kinase inhibition reduces the transport of IcMHCII to the cell surface to engage in antigen presentation. Indeed, it was observed that, under certain conditions, perinuclear clustering of IcMHCII could be observed (Fig. 4A, C). When the percentage of cells exhibiting this perinuclear IcMHCII was quantified, it was observed that *G2019S* expressing pMacs had a significant decrease in cells with perinuclear IcMHCII relative to *WTOE* and B6 pMacs, with a significant decrease also observed in WTOE relative to B6 pMacs (Fig. 4C). Furthermore, PF360 treatment significantly increased the percentage of cells with perinuclear icMHCII, further supporting a role of Lrrk2 in transport of MHCII to the cell surface. A similar effect of LRRK2 ASO was also observed (Fig.4E).

**Figure 4.**
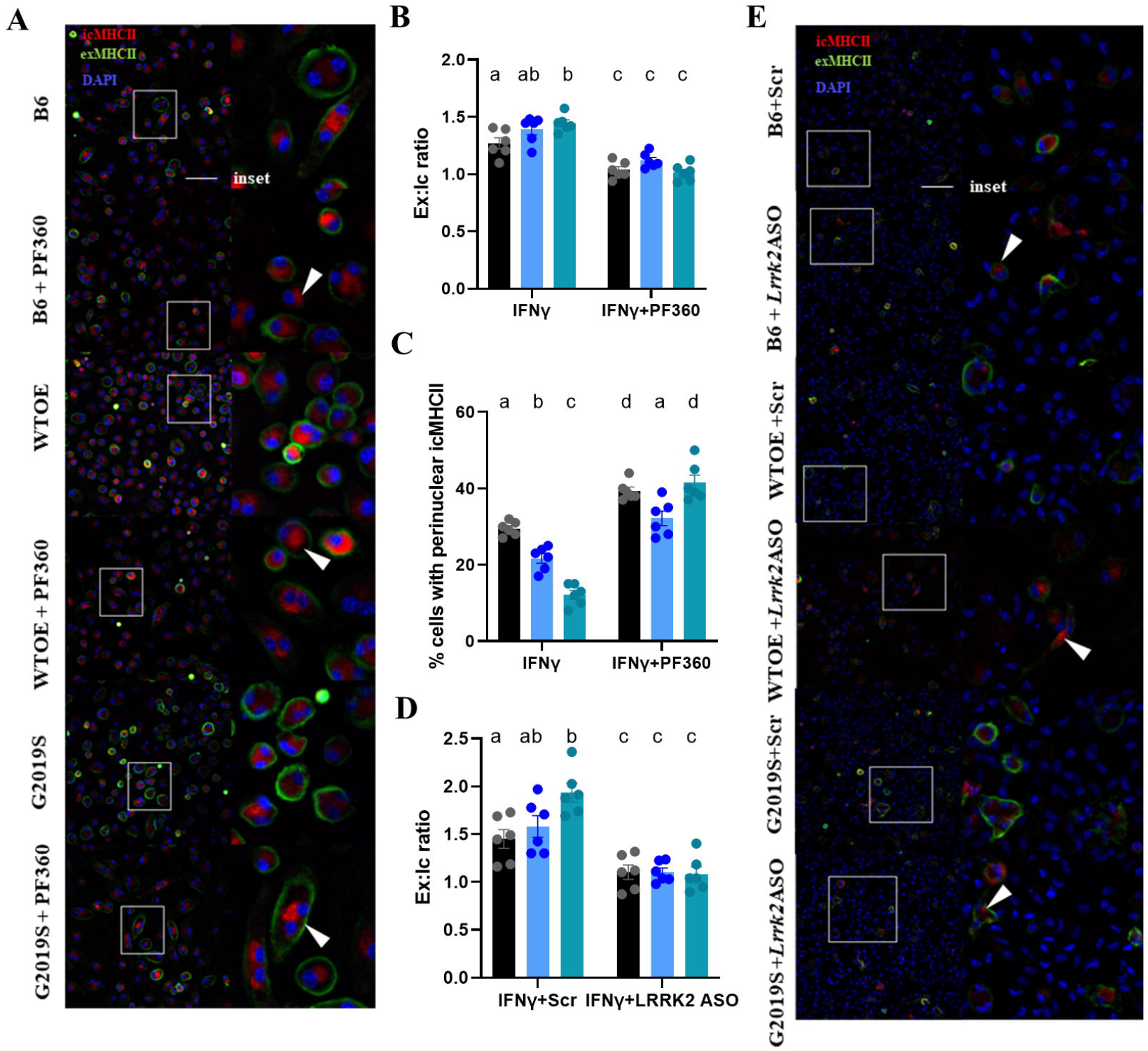
Lrrk2 kinase inhibition and knock-down via ASO reduces trafficking of MHC-II to the plasma membrane. pMacs from 10-12-week-old male B6, *WTOE* or *G2019S* mice were stimulated with 100U IFNy for 18-hours and stained for intracellular and extracellular MHC-II MFI and ex:ic ratio quantified. White arrows indicate perinuclear clusering of icMHCII. Scale bars, 30μM (**A, B**). Percentage of cells with perinuclear clustering was quantified (**C**). pMacs from 10-12-week-old male B6, *WTOE* or *G2019S* mice were nuclefoected with a *Lrrk2*-targetting ASO or scramble control and stimulated with 100U IFNγ for 18-hours and stained for intracellular and extracellular MHC-II MFI and ex:ic ratio quantified. White arrows indicate perinuclear clusering of icMHCII. Scale bars, 30Μm (**D, E**). Bars represent mean +/- SEM (N = 6). Two-way ANOVA, Bonferroni post-hoc, groups sharing the same letters are not significantly different (p>0.05) whilst groups displaying the same letter are significantly different (p<0.05).

### *Lrrk2* knock-down via antisense oligonucleotide and kinase inhibition alters cytokine release from pMacs

To see if alterations in antigen presentation and lysosomal function in Lrrk2 ASO or Lrrk2 kinase inhibitor treated cells were accompanied by changes in cytokine release, media from vehicle-and IFNγ treated cells were collected and cytokine levels quantified. It was observed that both ASO-mediated Lrrk2 knock-down and Lrrk2 kinase inhibition was able to significantly reduce IFNγ-dependent IL10 release in *G2019S-* expressing pMacs (Fig.5A,B), suggesting IL10 release from stimulated pMacs may be dependent on Lrrk2, specifically its kinase activity. This same observation was made regarding TNF release from *G2019S* pMacs (Fig5C,D). A significant reduction in IL12p70 release in *G2019S* pMacs nucleofected with the *Lrrk2-* targetting ASO (Fig. 5E). On the other hand, although ASO-mediated Lrrk2 knock-down was capable of decreasing IL4 release in IFNγ-treated *G2019S*-expressing pMacs (Fig.5F), Lrrk2 kinase inhibition was unable to do so (Sup.Fig.5A). No significant effects of ASO treatment or Lrrk2 kinase inhibition on cytokine release were observed in other cytokines measured (Sup.Fig5B-E).

**Figure 5.**
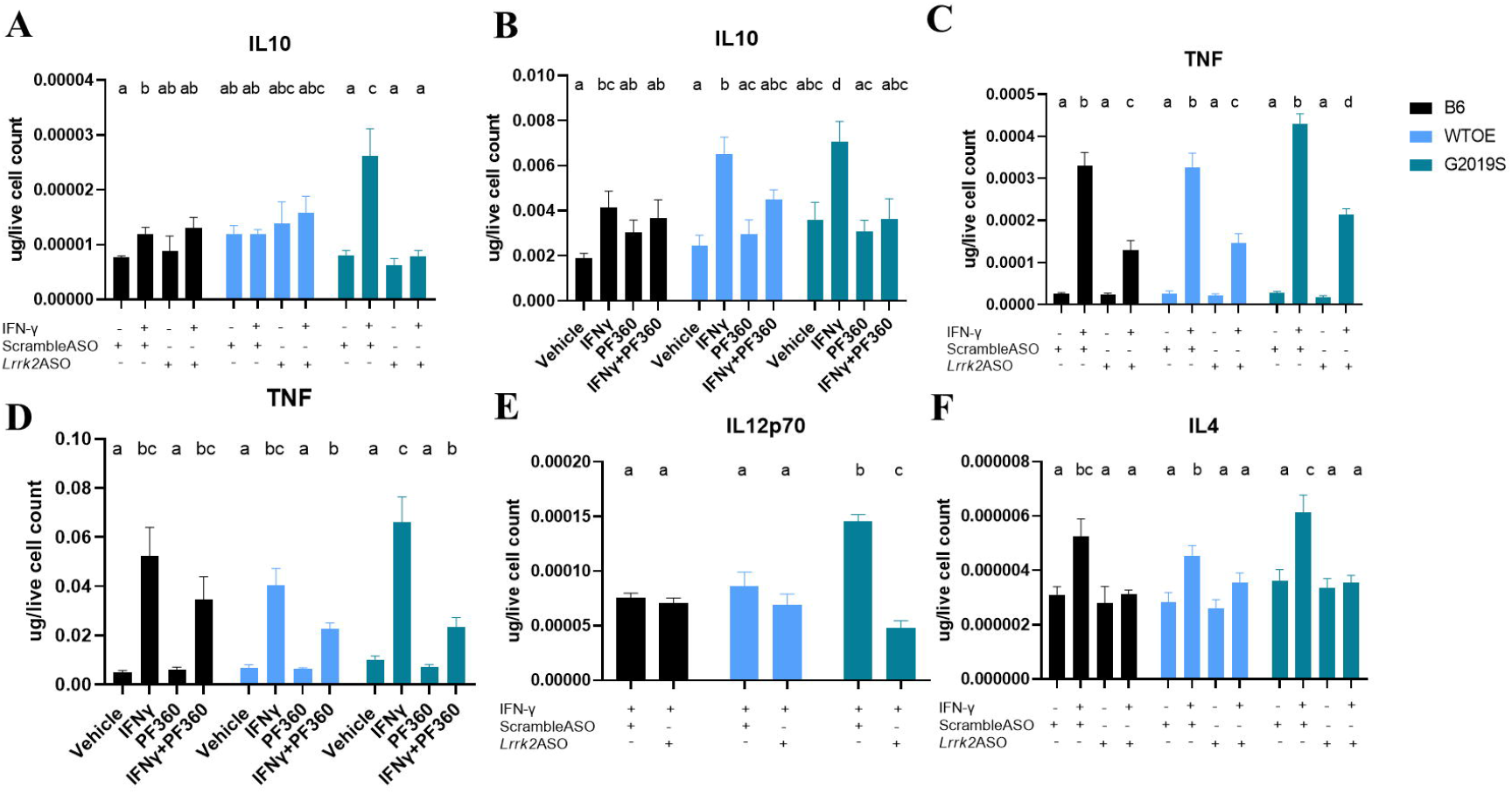
LRRK2 knock-down via antisense oligonucleotide and kinase inhibition alters cytokine release from pMacs: pMacs from 10-12-week-old male B6, *WTOE* or *G2019S* mice were nuclefoected with a *Lrrk2*-targetting ASO or scramble control and stimulated with 100U IFNγ, or were plated with 100U IFNγ +/- 100nM of Pf360 and media collected after 18-hours. Cytokine levels of IL10 (**A, B**), TNF (**C, D**), IL12p70 (**E**) and IL4 (**F**) were quantified and normalized to live cell count. Bars represent mean +/- SEM (N = 8-10). Two-way ANOVA, Bonferroni post-hoc, groups sharing the same letters are not significantly different (p>0.05) whilst groups displaying the same letter are significantly different (p<0.05).

### ASO-mediated Lrrk2 knock-down alters critical immune pathways in *G2019S* and *WTOE* pMacs

*Lrrk2* mRNA was significantly reduced in WTOE and G2019S pMacs in both vehicle and IFNγ treated conditions (Fig. 6A). It is also interesting to note that in G2019S pMacs nucleofected with the Lrrk2 ASO, a significant reduction in Lrrk2 mRNA is observed in response to IFNγ, as opposed to the increase in Lrrk2 expression previously reported in response to inflammatory stimulus.

**Figure 6.**
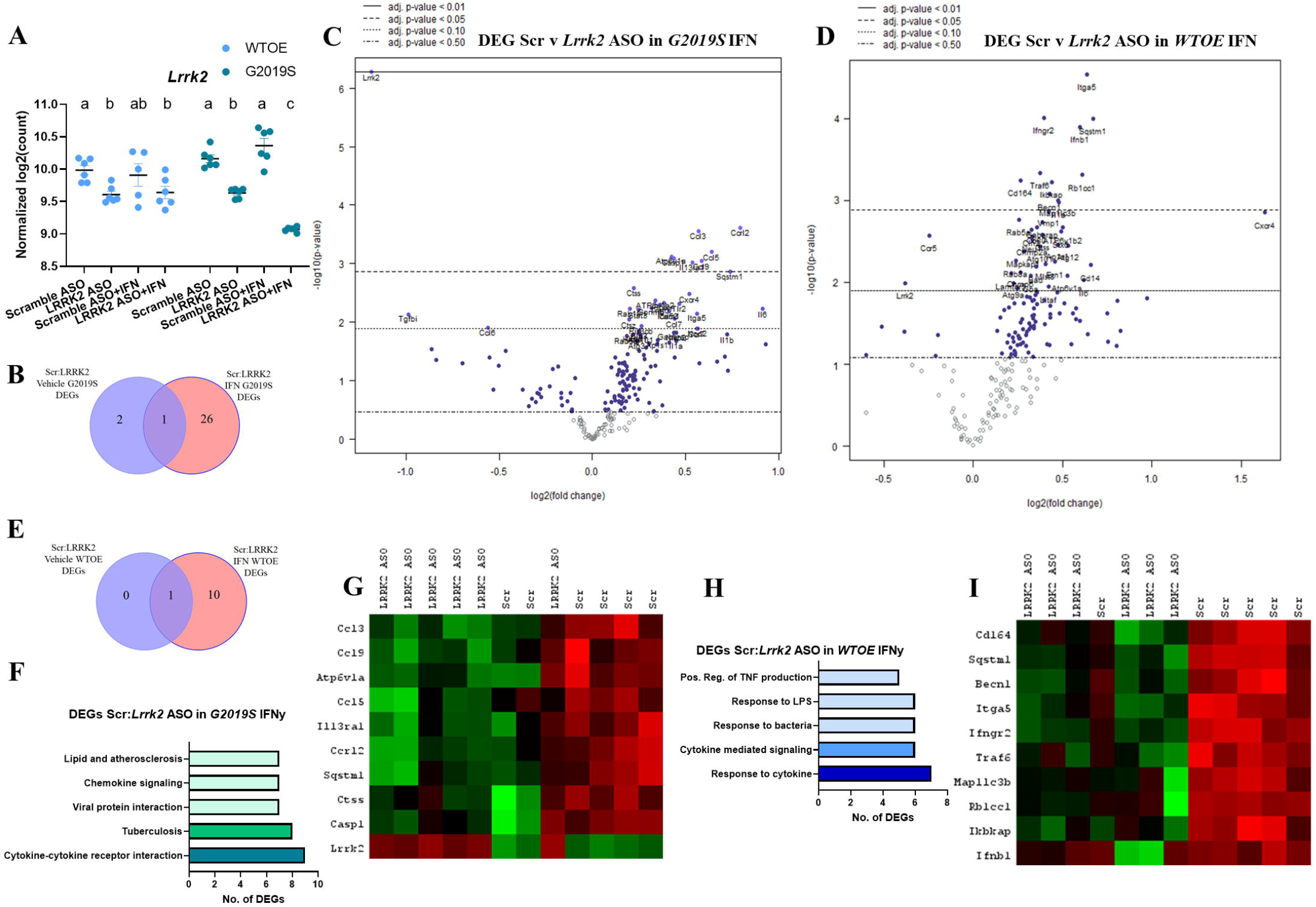
ASO-mediated Lrrk2 knock-down alters critical autophagy and cytokine signalling pathways in *G2019S* and WTOE pMacs: Transcriptomic analysis from *WTOE* or *G2019S* vehicle and IFNγ-treated pMacs nucleofected with scramble or *Lrrk2* ASO. (**A**) Normalized log2(count) value for *Lrrk2* mRNA was plotted. Bars represent mean +/- SEM (N = 6). Two-way ANOVA, Bonferroni post-hoc, groups sharing the same letters are not significantly different (p>0.05) whilst groups displaying the same letter are significantly different (p<0.05). (**B, C**) Significant Scramble: *Lrrk2* DEGs in *G2019S* pMacs were counted and compared across vehicle and IFNγ treatments. (**D, E**) Significant Scramble: *Lrrk2* DEGs in WTOE pMacs were counted and compared across vehicle and IFNγ treatments. Volcano plots show proteins with fold change□>□1.5 and an adjusted p-value□≤□0.05. (**F**) ShinyGO 0.76.3 was used to identify pathways in which significant DEGs were associated with. (**G**) Heat maps show DEGs seen only in *G2019S* pMacs treated with IFNγ. (**H**) ShinyGO 0.76.3 was used to identify pathways in which significant DEGs were associated with. (**I**) Heat maps show DEGs seen only in WTOE pMacs treated with IFNγ.

It was observed that an increase in number of scr:LRRK2 DEGs was observed in IFNγ treated *G2019S* pMacs relative to those seen in vehicle treated conditions (Fig. 6B, C). A similar observation was observed in *WTOE* pMacs (Fig. 6D, E). Given that Lrrk2 protein levels increase in response to IFNγ, it follows that in conditions where Lrrk2 may play a more crucial and fundamental role in immune cells that knock-down of Lrrk2 would have a more significant effect.

Using ShinyGo analysis to identify pathways in which these DEGs were situated in, the scramble: *Lrrk2* DEGs identified in *G2019S* and *WTOE* pMacs treated with IFNγ were found in critical immune-related pathways such as cytokine-cytokine receptor interaction, TNF production and responses to bacteria (Fig. 6F-I). It was also noted that critical autophagy-related genes, such as *sqstm1, ctss, becn1* and *Atp6v1a* were up-regulated when Lrrk2 levels were decreased as it has previously been described that mTOR inhibition, and therefore increased expression of autophagy-related genes and increased autophagic flux, decreases antigen presenting capabilities of AP cells [32].

### Nanostring-based transcriptome analysis reveals genotype differences in a treatment specific manner and reveals differential response to IFN-γ by *G2019S* pMacs

To obtain insights into the role of Lrrk2 in lysosomal function during the inflammatory response, and the effects of Lrrk2 knock-down on this, we performed a NanoString-based mRNA expression analysis of selected lysosomal and autophagy related genes in pMacs nucleofected with scramble or *Lrrk2* ASO, treated with vehicle or IFNγ for 18-hours.

Firstly, we wanted to identify key cellular pathways that were altered in *G2019S* pMacs relative to *WTOE* pMacs in scramble ASO conditions, in both vehicle and IFNγ-treated conditions; that is, the effects of increased kinase activity levels in a context-dependent manner. Although a degree of overlap was seen regarding WTOE:G2019S DEGs identified in both vehicle and IFNγ treatments, a number of DEGs were identified as novel to each of these treatment conditions (Sup.Fig. 6A-D). ShinyGO pathway analysis showed that WTOE:G2019S DEGs seen in both treatment conditions were identified in pathways such as glycosylceramide catabolic processing, lysosomal pH, and lysosomal/vacuole organization (Sup.Fig. 6E). A number of these DEGs have previously been identified as substrates or interactors of Lrrk2 [33–35] and have previously been implicated in PD [36–38] (Sup.Fig. 6F). These DEGs were downregulated in *G2019S* pMacs relative to *WTOE* pMacs, indicating a potential down regulation of these pathways in *G2019S* pMacs. WTOE:G2019S DEGs seen in vehicle treatment only were identifed in pathways related to cytokine production, signal transduction, and cellular communication (Sup.Fig. 6G,H). Interestingly, WTOE:G2019S DEGs seen in IFNγ treatment only were most identified in pathways associated with vesicle transport, and Rab and Ras signal transduction (Sup.Fig. 6I, J). These DEGs were downregulated in *G2019S* pMacs relative to *WTOE* pMacs, indicating a potential down regulation of these pathways in *G2019S* pMacs. Again, many of these DEGs have previously been identified as substrates of Lrrk2 [35] and have previously been implicated in PD as well as other neurodegenerative diseases [39–41].

### Nanostring-based transcriptome analysis reveals differential response to IFN-γ by G2019S pMacs

We were also interested to understand how increased Lrrk2 kinase activity may modulate macrophages responses to an inflammatory stimulus, and if vehicle:IFNγ DEGs in *G2019S* pMacs differed from those in *WTOE* pMacs. There was a vast degree of overlap between vehicle:IFNγ DEGs in the two genotypes, however, 32 DEGs were identified in *G2019S* pMacs that were not found in *WTOE* pMacs (Sup.Fig. 7A-D). These 32 DEGs were identified in pathways related to autophagy, immune cell activation and regulation of phosphorylation (Sup.Fig. 7E). Most interestingly, the pathway termed ‘regulator of neuronal death’ was significantly enriched by these DEGs. DEGs identified in this pathway include *Lgmn, Bad, Casp2, Tlr4, Grn, IL18, Cln3* and *Gba*, with the majority of these genes being downregulated upon IFNγ treatment in *G2019S* pMacs (Sup.Fig. 7F).

### Vesicular trafficking and lysosomal positioning pathways are associated with the response to IFNγ in *G2019S* pMacs nucloefected with *Lrrk2* ASO

We also wanted to understand how *Lrrk2* knock-down affects the response to IFNγ in G2019S pMacs, and begin to unveil a mechanism of action regarding the capabilities of *Lrrk2* ASO treatment to ameliorate antigen presentation and lysosomal phenotypes observed in *G2019S* pMacs here. When comparing Vehicle:IFNγ DEGs in scramble and *Lrrk2* ASO treated *G2019S* pMacs, we found that, although many of the DEGs persisted in the *Lrrk2* ASO condition, suggesting no effect of *Lrrk2* knock-down on these DEGs, 15 DEGs were novel to scramble conditions and 34 DEGs were seen only in Lrrk2 ASO treated cells (Fig.7A-C). Interestingly, a number of the Vehicle:IFN γ DEGs in scramble ASO - treated G2019S pMacs only included those that enriched the ‘regulator of neuronal death’ pathway previously described (Fig.7D). The fact that these DEGs are only seen in scramble conditions and not in Lrrk2 ASO conditions suggests that the knock-down of *Lrrk2* sufficiently prevents these changes in gene expression in response to IFNγ in *G2019S* pMacs. Interestingly, Vehicle:IFNγ DEGs identified in *Lrrk2* ASO-treated G2019S pMacs were identifed in pathways related to acute inflammatory responses to antigen, lysosome localization and exocytosis (Fig. 7E). A number of the genes identified in exocytosis and lysosome positioning pathways have specifically been identified to play a role in the release of secretory lysosomes, exocytosis of lysosomes and transportation of lysosomal content to the plasma membrane, including *Snap23, Abca1, Tmem55b, Mtor, Rab3a and Rab3d* [21, 42–46] (Fig.7F), and these were all down-regulated in response to IFNγ in *G2019S* pMacs treated with Lrrk2 ASO. Interestingly, LRRK2 has recently been shown to mediate tubulation and vesicle sorting from lysosomes [20, 47]. with *G2019S-Lrrk2* expression significantly increasing the formation of these tubules from lysosomes. In the context of myeloid cells, it is known that lysosomal tubulation is usually observed in cells undergoing immune activation, and these tubules are crucial for both; phagocytosis and antigen presentation [32, 48]. Collectively, therefore, we hypothesized that increased lysosomal activity and antigen presentation observed in *G2019S* pMacs here may be due to increased LTF, and that knock-down of Lrrk2 with ASO may ameliorate these phenotypes by reducing LTF.

**Figure 7.**
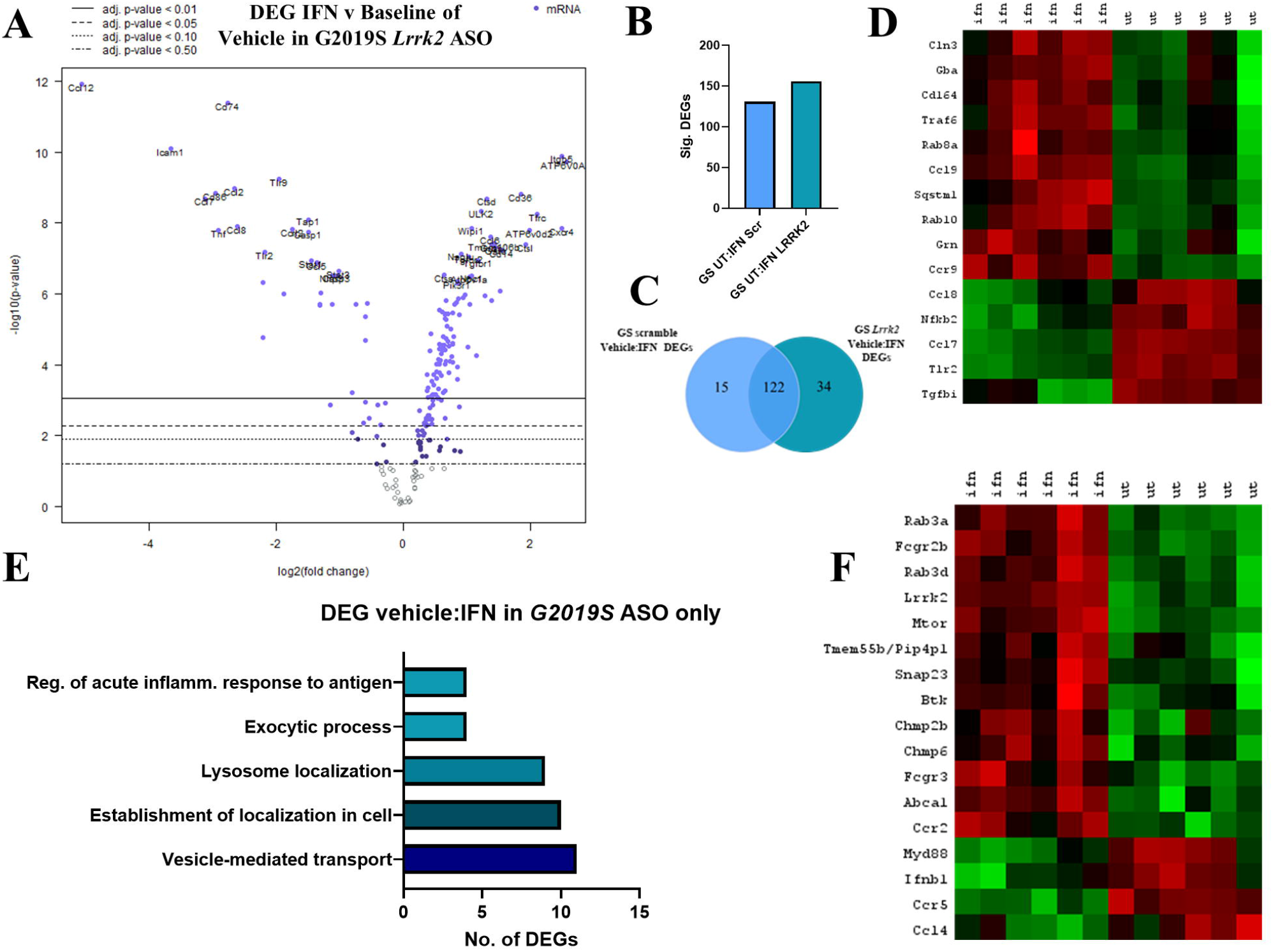
Vesicular trafficking and lysosomal positioning pathways are associated with the response to IFNγ in *G2019S*pMacs nucloefected with *Lrrk2* ASO: Transcriptomic analysis from *Lrrk2*-ASO nucleofected *G2019S* pMacs treated with vehicle or IFNγ (**A**). Volcano plot shows proteins with fold change□>□1.5 and an adjusted p-value□≤□0.05. (**B, C**) Significant DEGs were counted and compared across genotypes. (**D**) Heat maps show DEGs seen only in Scramble ASO nucleofected *G2019S* pMacs. (**E**) ShinyGO 0.76.3 was used to identify pathways in which significant DEGs were associated with. (**F**) Heat maps show DEGs seen only in *G2019S* pMacs nucleofected with *Lrrk2-* ASO.

### LRRK2 modulates antigen presentation via lysosomal tubule formation

In order to determine if alterations in antigen presentation in *G2019S* pMacs were due to altered LTF, pMacs were loaded with Dextran-Alexa Fluor 546™ for 1 hour, followed by a 2-hour pulse period to ensure Dextran was fully loaded into lysosomal compartments and treated with 100U of IFNγ to induce LTF. Live cells were imaged after 2 hours at which LTF would have occurred [42]. When quantifying the percentage of cells with lysosomal tubules (defined as >2µM), it was observed that IFNγ-treated *G2019S* pMacs exhibited significantly increased percentage of cells with tubular lysosomes relative to B6 controls and *WTOE* pMacs (Fig8A, B, C; Sup. Fig7A). It is known that tubular lysosomes that form in immune cells for the purpose of phagocytosis and antigen presentation are dependent on mTOR activity, whereas tubular lysosomes that form for the purpose of autophagy are not [49]. Therefore, to differentiate between these two functions, cells were co-treated with the mTOR inhibitor, Tornin1, a significant reduction in cells with tubular lysosomes was observed in all genotypes (Fig8B). When this was quantified as a percentage of cells with Torin1-dependent tubular lysosomes, the significant increase in *G2019S* pMacs relative to other genotypes persisted (Sup.Fig7B), suggesting that the tubular lysosomes quantified here, and those that are increased in *G2019S* pMacs are mTOR-dependent and therefore involved in antigen presentation. When cells were co-treated with the LRRK2 kinase inhibitor, PF360, a significant reduction was also observed in all genotypes, suggesting LTF in macrophages in dependent on LRRK2 kinase activity. Similarly, when cells were nucleofected with a *Lrrk2* ASO, a significant reduction in LTF was observed relative to scramble controls (Fig8C; Sup.Fig7C).

**Figure 8.**
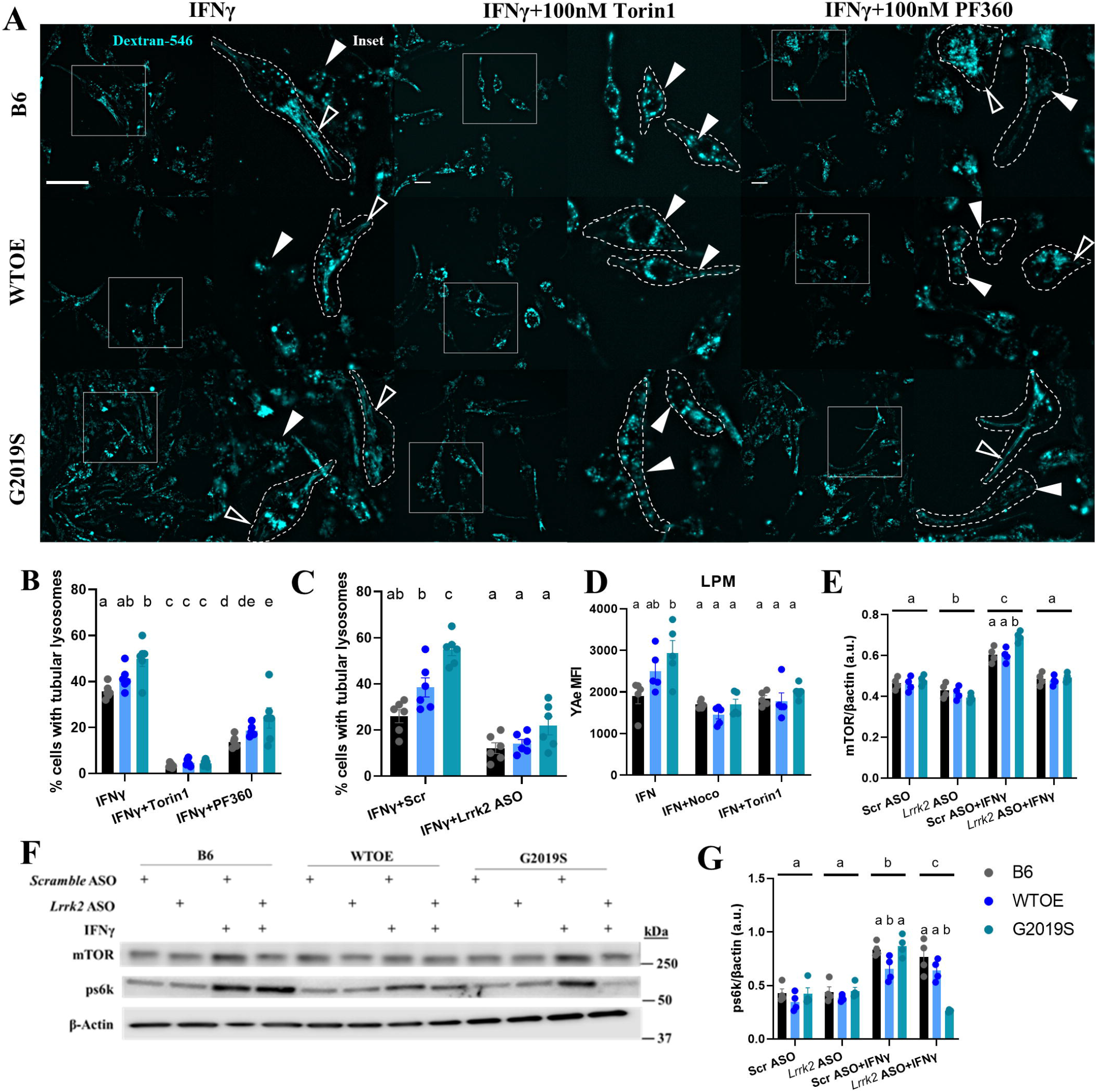
LRRK2 modulates antigen presentation via lysosomal tubule formation. pMacs from 10-12-week-old male B6, WTOE or G2019S mice treated with 0.5mg/mL Dextran Alexa-Fluor546 for 1-hour, followed by a 2-hour pulse-period to ensure loading into lysosomes, and then treated with 100U of IFNy +/- 100nM Torin1 or 100nM PF360 for 2-hours to stimulate LFT. (**A**) Cells were imaged live, and (**B**) percentage of cells with tubular lysosomes quantified. Filled white arrows indicate pMacs with tubular structures, empty arrow heads indicate pMacs with punctate dextran. Dotted lines indicate masks of cells based on bright field images. Scale bars, 10μM (**C**) Cells were nucleofected with 1uG of scramble of *Lrrk2*-targetting ASO and percentage of cells with tubular lysosomes quantified. (**D**) pMacs were treated with 100U of IFNy +/- 10uM Nocodazole or 100nM Torin1 for 18-hours and YAe MFI in LPMs was quantified via flow-cytometry. (**E, F, G**) pMacs were treated with 100U of IFNy +/- 100nM PF360 and protein lysate assessed for mTOR, and s60k protein levels and normalized to β-actin levels and quantified. Representative western blots shown. Bars represent mean +/- SEM (N = 4-6). Two-way ANOVA, Bonferroni post-hoc, groups sharing the same letters are not significantly different (p>0.05) whilst groups displaying the same letter are significantly different (p<0.05).

Fluorescently labeled dextrans have been shown to enter the cell by both macro-[50] and micropinocytosis [51]. It has recently been demonstrated that LRRK2 and Rab10 coordinate micropinocytosis in human and mouse phagocytic cells [52]. In order to ensure that differences in dextran-labelled tubules were not due to differences in uptake between genotypes in our study, cells were loaded with 20μg of dextran for 1 hour, fixed and imaged with no pulse phase and dextran-fluorescence quantified. As well, to ensure other endocytic pathways which may affect uptake of dextran [53] are not significantly different between genotypes, cells were incubated with Alexa Fluor594-transferrin and uptake measured via fluorescent microscopy. No significant effects of genotype or LRRK2 kinase inhibition on transferrin or dextran uptake were observed (Sup.Fig.7D-G), although a significant reduction in transferrin uptake was observed upon IFNγ treatment.

To further explore the role of LTF in the *G2019S*-associated phenotypes observed here, we repeated the Yae antigen-presentation assay described before with co-treatments with compounds known to inhibit various steps of the antigen presenting process that depends on LTF. Both the previously described Lrrk2-mediated tubulation [20] and antigen presentation via tubules are dependent on microtubules [54]; co-treatment of IFNγ-treated pMacs with the microtubule destabilizer, nocodazole, significantly reduced YAe MFI in LPMs, indicating decreased antigen presenting abilities of these cells, in all three genotypes and ameliorating the *G2019S*-dependent increase observed in vehicle treated cells (Fig.8D). Such observations suggest that the *G2019S*-dependent increase in antigen presentation in LPMs is dependent on microtubules. mTOR inhibition has also been shown to decrease LTF-dependent antigen presentation in dendritic cells and macrophages [32]. Treatment with the mTOR inhibitor, Torin1, also decreased Yae MFI in LPMs in all genotypes, ameliorating any effects of the *G2019S* mutation relative to the other two genotypes (Fig.8D). As previously discussed, presentation and pathogen sensing requires protease action and sufficient lysosomal function in order to occur [30], and treatment with the vATPase-A1 pump inhibitor, Bafilomycin A1, significantly reduced YAe MFI in these cells (Fig.1H).

We next wanted to determine if the trafficking of MHC-II to the plasma membrane was dependent on mTOR-dependent LTF. We found that Torin1 co-treatment induced the same effects as Lrrk2-kinase inhibition and knock-down on Ex:Ic MHC-II ratio and nuclear clustering (Sup.Fig.9A, B, C). This suggests that the LTF-dependent MHC-II antigen presentation phenotype observed is mTOR-dependent. This is further supported by the fact that mTOR was a significant DEG observed in the *Lrrk2* ASO nanostring-based transcriptomics in *G2019S* pMacs (Fig.7F), with mTOR expression decreasing in response to IFNγ only in cells treated with the *Lrrk2* ASO.

Inhibition of mTOR is a key trigger for autophagy. Therefore, we wondered whether the observed decrease in LTF in cells treated with *Lrrk2* kinase inhibitor was accompanied by an up-regulation in autophagic flux. Furthermore, we hypothesized that the increased stimulation-dependent LFT observed in *G2019S* cells would be accompanied by a decrease in autophagic flux. Indeed, when quantifying LC3-II flux (calculated as signal difference between conditions with and without Bafilomycin A1), it was observed that all genotypes exhibited decreased autophagic flux upon IFNγ-treatment, with this effect exacerbated in *G2019S* pMacs (Sup.Fig9.D,E). Treatment with either the LRRK2 kinase inhibitor, PF360, or the mTOR inhibitor, Torin1, ameliorated this stimulation-dependent effect on LC3 flux in all genotypes. Collectively such data suggests that there is a delicate balance between prioritizing autophagic flux or LTF in macrophages, and that both Lrrk2 and mTOR are key components in regulating and maintaining this balance.

Given this, we wanted to assess whether *G2019S* pMacs exhibited an increase in mTOR expression and/or activity levels which may be driving LTF in these cells. Indeed, we saw a significant increase in mTOR protein levels in *G2019S* pMacs in a stimulation-dependent manner, and this was ameliorated with the knock-down of *Lrrk2* via ASO (Fig.8E, F). Furthermore, we saw a significant increase in phosphorylated s6k levels (p70S6 kinase 1), a downstream target of mTOR signaling, in *G2019S* pMacs upon IFN γ stimulation which was ameliorated with the knock-down of Lrrk2 via ASO (Fig. 8F, G). As well, we observed an increase in RILPL1 (Rab interacting lysosomal protein like 1 expression), a modulator of lysosomal positioning and a previously identified interactor of LRRK2 [55], in *G2019S* pMacs upon IFN γ stimulation, which was ameliorated with the knock-down of Lrrk2 via ASO (Sup.Fig.9F, G). Collectively, this data suggests that the knock-down of *Lrrk2* in *G2019S* pMacs ameliorates LTF-dependent antigen presentation by modulating mTOR levels and activity.

## Discussion

Recent advances in understanding LRRK2 function at the lysosome have suggested a potential link for the role of LRRK2 on the regulation of lysosomal function to that of immune cell function and modulation of inflammatory responses. Our data demonstrate increased antigen presentation, cytokine release, and lysosomal activity in pMacs from *G2019S* mice which were successfully ameliorated with knock-down of *Lrrk2* via ASO or treatment with a LRRK2 kinase-inhibitor. Our findings suggest that increased antigen presentation in mutant *Lrrk2* cells is intrinsically linked to the alterations in lysosomal function observed, with inhibition of lysosomal function via Bafilomycin A1 causing decreased antigen presentation in these cells. Our LRRK2 kinase inhibitor findings also demonstrate that the role of Lrrk2 in antigen presentation occurs early on in the response to IFNγ, and is accompanied by alterations in lysosomal peak activity during this response. Furthermore, the effects of Lrrk2 appear to be specifically mediated through its kinase activity. It was discovered that increased LTF is the underlying mechanism for the phenotypes observed in *G2019S* pMacs, in a potentially mTOR-dependent manner, with *Lrrk2* ASO and kinase inhibition ameliorating this.

We found that release of three cytokines were consistently elevated in *G2019S* pMac cultures: IL10, IL4 and TNF. It has recently been shown that RAW264.7 cells expressing *T1348N-LRRK2*, an artificial P-Loop null mutation that disrupts GTP binding [56, 57], produce significantly less IL10 relative to wild-type cells in response to LPS and Zymosan [58]. IL10 is produced by macrophages and is critical in limiting immune-mediated pathology [59], and it was therefore suggested by Nazish et al that there may be a neuroprotective role of LRRK2 in immune signalling through altered IL10 secretion. Indeed, IL4, which is also critical in resolving inflammation [60], was increased in *G2019S* pMac cultures in this study. Although a pro-inflammatory cytokine, the fact that TNF was also increased in *G2019S* pMac cultures in this study supports the hypothesis of LRRK2 being protective in the immune system given the fact that mice lacking TNF and/or TNF receptors have been reported to exhibit increased susceptibility to infection with increased bacterial load and increased inducible nitric oxide species (iNOS) production [61]. In agreement with the hypothesis that LRRK2 may play a protective role in immune signalling, Lrrk2 has been shown to be required for efficient control of certain pathogens; LRRK2 has been implicated in the control of the enteric pathogen *Salmonella typhimurium* via NLRC4 inflammasome regulation in macrophages from Lrrk2-KO mice [14, 15]. This is supported by the observation that *G2019S* knock-in mice, controlled *Salmonella typhimurium* infection better, with reduced bacterial growth and longer survival during sepsis; an effect which was dependent on myeloid cells [62]. Furthermore, paneth cells from *Lrrk2*-KO mice are more susceptible to infection from *Listeria monocytogenes*, with a loss of Lrrk2 causing decreased levels of lysozyme, an antimicrobial enzyme responsible for the degradation and lysis of bacteria [16]. It is possible that increased LTF and therefore antigen presentation in *G2019S* pMacs is the underlying mechanism of the role of Lrrk2 in pathogen control and its protective role in the immune system. However, other reports describe a deleterious role of Lrrk2 in pathogen control. For example, animals with reovirus-induced encephalitis that expressed the *G2019S* mutation exhibited increased mortality, increased reactive oxygen species and higher concentrations of α-synuclein in the brain [62]. Furthermore, loss of *LRRK2* enhances *Mycobacterium tuberculosis* (Mtb) control and decreases bacterial burdens in both primary mouse macrophages and human iPSC□derived macrophages [63]. It seems therefore that the role of Lrrk2 in pathogen control may be pathogen specific as well as cell-and tissue-type specific. *Salmonella typhimurium* and *Listeria monocytogenes* are both food-borne pathogens that enter the body through the gut, whereas reovirus and Mtb are air-borne that enter through the lungs. Indeed, macrophages in the body are heterogeneous, showing specific transcription factors and markers and therefore different functions in the body [64]. It is possible therefore that the role Lrrk2 plays is different between different macrophages subtypes, and may also be sex- and age-dependent [62, 65, 66]. In summary, we have described a role of LRRK2 in LTF which can be disrupted by ASO knock-down, however further research is required to determine if this is seen in other antigen presenting cell types and whether such phenotypes alter pathogen control in these cells.

The use of both kinase inhibitors and knock-down of total Lrrk2 in this study also sheds light on the role of the kinase domain vs other enzymatic and protein-protein interaction domains of Lrrk2. For example, it was observed that both 18-hour Lrrk2 kinase-inhibition and *Lrrk2* ASO treatment were capable of ameliorating *G2019S*-associated increases in antigen presentation and LTF, suggesting a role of the Lrrk2 kinase domain in these functions. Lrrk2 has recently been implicated in a function termed LYsosomal Tubulation/sorting driven by LRRK2, or LYTL [20]. Bonet-Ponce and colleagues reported that LRRK2 recruits JIP4 to lysosomes in a kinase-dependent manner via the phosphorylation of RAB35 and RAB10, promoting the formation of tubules in response to lysosomal membrane damage. Rab35 was indeed identified in the Nanostring nCounter analysis in this study with Rab35 mRNA being down-regulated in *G2019S* IFNγ-treated pMacs relative to *WTOE*. Such a down-regulation may be a compensatory mechanism for increased Rab35 phosphorylation. Additional research is needed to determine if similar Lrrk2 substrates are involved in LYTL and LTF; although similar, tubular lysosomes are known to play a role in multiple functions and therefore different interacting partners and characteristics of these tubular lysosomes are likely [47].

It is interesting that, although IL10 and TNF production was decreased by Lrrk2 kinase-inhibition in this study, IL4 release was unaffected by Lrrk2 kinase-inhibition and was only ameliorated upon knock-down of Lrrk2, suggesting a kinase-independent role of Lrrk2 in the release of IL4 from pMacs. Cytokine secretion pathways are often adapted to suit specific cytokines, their function, and cell type. Macrophages lack granules, which enable rapid release of the cytokines upon cell activation, and instead cytokines must be synthesized after cell activation and secreted [67]. Three canonical transport pathways for cytokine secretion have been identified to date; direct transport to the cell surface from the trans Golgi network, via the recycling endosome, and during phagocytosis where cytokine is routed from the recycling endosome to the phagocytic cup [67]. Notably, TNF and IL10 have been detailed to be processed and released via these canonical pathways, whereas much less is known about the non-classical secretory pathways for cytokine release thought to be utilized by cytokines such as IL1β, IL18 and IL4 [67]. It is therefore possible that Lrrk2 mediates these canonical pathways via kinase-activity, which is plausible given the numerous reports identifying Rabs, known for regulating protein transport, vesicle trafficking and membrane fusion, as *bona fide* kinase substrates of LRRK2 [35, 68]. Additional research is required to understand the role of LRRK2 in non-canonical pathways of cytokine release and the requirement of LRRK2 GTPase activity and protein-protein interaction domains in this role.

LRRK2 has previously been shown to coordinate macropinocytosis via Rab10-recruitment to macropinosomes, which are MHC-II and Cd11b positive, and induce subsequent CL5- stimulated Akt signalling and bone marrow-derived macrophage chemotaxis [52]. Whilst Lrrk2-kinase inhibition decreased chemotaxis in these cells, increased surface receptor recycling was seen with Lrrk2 kinase-inhibition. It has previously been described that mature macropinosomes will fuse with tubular lysosomes that mediate their contents to the cell surface [52]. It may be, therefore, that LRRK2 plays multiple roles in the processing and trafficking of MHC-II to the cell surface to engage in antigen presentation, thus the role of LRRK2 in macropinocytosis could be a potential confound in this study with multiple interpretations of the data. For example, increased MHC-II expression on LPMs in this study may be due to increased surface receptor recycling as opposed to increased antigen presentation due to increased LTF; however, Liu et al observed an increase in surface receptor recycling during Lrrk2 kinase-inhibition, so therefore we would not expect this mechanism to be involved in *G2019S*pMacs which exhibit increased kinase activity. Furthermore, Liu et al 2020 looked at MHC-II expression, as opposed to MHC-II complexes loaded with a peptide for antigen presentation; the use of the YAe assays in this study and the fact we can quantify antigen presenting MHC-II complexes leads us to conclude that the mechanism we are measuring here is indeed antigen presentation via LTF. Furthermore, no differences were observed in the uptake levels of Dextran or transferrin via micropinocytosis or receptor-mediated endocytosis between genotypes, with no effects of Lrrk2 kinase inhibition, suggesting these mechanisms are unlikely playing a role in the phenotypes observed. However, it will still be of interest to future studies to unveil how macropinocytosis and LTF for antigen presentation interact and how LRRK2 may be implicated in this.

Interestingly, we observed different pathways altered and DEGs between *WTOE* and *G2019S* pMacs depending on whether cells were stimulated with IFNγ or not. These data, and the fact that Lrrk2-kinase inhibitors had different effects on the phenotypes described here depending on when in the IFNγ-response it is administered supports the hypothesis that LRRK2 may behave as a ‘date-hub’. The ‘date-hub’ hypothesis describes two types of ‘hubs’, one of which are ‘date hubs’, which bind their different partners at different times or locations. The potential for LRRK2 behaving as a ‘date-hub’ has been discussed in the literature [69] and may explain the differences in DEGs and effects of kinase inhibition described here, and may also explain the discrepancies reported regarding conflicting results [14]. However, despite being discussed in the literature, no study has yet been completed to definitively show whether the LRRK2 interactome varies in a cell-type or time-dependent manner.

Intriguingly, *Rab8a, Rab8b, Rab35*, and *Rab5b* were identified as DEGs between *WTOE* and *G2019S* pMacs, only when treated with IFNγ. Such data is in agreement with previous literature that reports concomitant increases in LRRK2 and pRab10 in human monocytes upon IFNγ stimulation [13] and that LRRK2 is recruited to membranes upon LPS-stimulation [18, 19]. However, perhaps counterintuitively, mRNA of these Rabs are down-regulated in *G2019S* pMacs relative to *WTOE*. Further exploration is necessary regarding how these findings translate to the protein level for LRRK2 substrates. It may be, however, that mRNA levels are down-regulated in *G2019S* pMacs as a compensatory mechanism to compensate for increased phosphorylation of Rabs by Lrrk2 [35, 68].

It was noted that many of the DEGs identified were down-regulated in *G2019S* pMacs relative to *WTOE*, as well as DEGs identified in IFNγ relative to vehicle-treated *G2019S* cells. It may be, therefore, that in *G2019S* pMacs, in particular those treated with IFNγ, exhibit an overall downregulation of gene expression. TFEB is a master regulator of autophagy- and lysosome-related genes, is known to regulate cytokine production in immune cells [70], and is inhibited by mTOR activity [71]. Interestingly, mTOR is required for LPS-induced lysosome tubulation and presentation of MHC-II in macrophages and dendritic cells, with mTOR inhibition decreasing lysosomal tubulation [42]. It therefore follows that in this study, one potential mechanism of action for the downregulation of gene expression in *G2019S* pMacs in response to IFNγ may be increased mTOR activity, leading to increased TFEB inhibition and decreased transcription, with concomitant increases in LTF. In support of this mechanism, we found that *G2019S* pMacs exhibited increased mTOR levels and mTOR activity upon immune stimulation, leading to increased LTF and decreased autophagic flux, all of which could be ameliorated by inhibition of mTOR activity via Torin1 treatment.

It is known that mTOR inhibition is a key trigger for autophagy [72]. In healthy individuals, mTOR signalling is responsible for maintaining a balance between protein synthesis, autophagy, and nutrient usage and storage processes. This balance is crucial for the cell since its dysregulation leads to cancer, obesity, and diabetes [73]. The lysosome surface serves as a platform to assemble major signalling hubs like mTOR, as well as AMPK, GSK3 and the inflammasome; these molecular assemblies integrate and facilitate cross-talk between signals and ultimately enable responses such as autophagy, membrane repair and microbe clearance [74]. Here we have shown a phenotype present in *G2019S* pMacs which increases LTF whilst decreasing autophagic flux. Collectively, such observations suggest LRRK2 may be a modulator of lysosomal responses in immune cells, with increased LRRK2 kinase activity favouring the formation of lysosomal tubules, and therefore pathogen control, over other lysosome-associated functions such as autophagic flux.

ASOs have been suggested as a potential therapeutic for LRRK2-PD, with many hypothesizing that targeting the increased LRRK2 levels and kinase activity will be beneficial to pathology. Although ASO-mediated knock-down of Lrrk2 in this study did ameliorate the *G2019S* phenotypes described, it also decreased many of the functional readouts in both B6 and WTOE pMacs, and the consequences of these effects on key immune functions such as infection control will need to be considered carefully. We have already discussed that alterations in pathogen control need to be explored regarding Lrrk2 ASOs and LTF in *G2019S* pMacs, and it therefore follows that this could have significant consequences on the use of Lrrk2-targetting approaches for therapies. If LRRK2 plays a critical role in pathogen and infection control in a pathogen or tissue-specific manner, it may mean that LRRK2- targetting therapies may need to be limited to mutant LRRK2 carriers and/or may need to be delivered in a compartment-targeted manner as opposed to systemically to avoid deleterious effects in tissues and cell-types where LRRK2 is crucial for immune responses.

## Materials and methods

### Animals

Bacterial artificial chromosome (BAC) transgenic mouse strains overexpressing either mouse mutant *G2019S-Lrrk2* (*G2019S*) or mouse wild-type *Lrrk2* (*WTOE*) have previously been characterised (Li et al., 2009, Yue et al., 2015) and were maintained in the McKnight Brain Institute vivarium (University of Florida) at 22°C at 60-70% humidity and animals were kept in a 12-hour light/dark cycle. C57BL/6 littermate controls were used for all studies, with *G2019S/WTOE* and C57BL/6 controls cohoused. All animal procedures were approved by the University of Florida Institutional Animal Care and Use Committee and were in accordance with the National Institute of Health Guide for the Care and Use of Laboratory Animals (NIH Publications No. 80-23) revised 1996. Male mice were aged to 8-10-weeks old and sacrificed via cervical dislocation.

### Harvesting and culturing of peritoneal macrophages and ex-vivo stimulation of non-nucleofected cells

Peritoneal macrophages were harvested from mice which had received a 1 ml intraperitoneal administration of 3% Brewer thioglycollate broth 72-hours prior collection. Mice also received Buprenorphine Sustained-Release every 48-hours for pain relief. Mice were sacrificed via cervical dislocation and abdomen sprayed with 70% Ethanol. Skin of the abdomen was split along the midline, taking care to avoid puncturing or cutting the abdominal cavity. 10 mL of cold RPMI media (Gibco; 11875119) was injected into the peritoneal cavity using a 27G needle. After gentle massaging of the peritoneal cavity, as much fluid was withdrawn as possible from the peritoneal cavity using a 25G needle and 10 mL syringe. Aspirated fluid was passed through a 70uM nylon filter onto 50mL falcon and pre-wet with 5mL of HBSS^-/-^. Filters were then washed twice with 5mL of HBSS^-/-^ and then tubes spun at 400 × *g* for 5 minutes at 4°C. Supernatant was aspirated and pellet resuspended in 3 mL pre-warmed growth media (RPMI, 10% FBS, 1% Pen-Strep). Cells were counted and viability recorded using trypan-blue exclusion on an automated cell-counter (Countess™; Thermo). Volume growth media was adjusted so that cells were plated at 5 × 10^5^/mL in 6-, 24- or 96-well plates depending on the intended assay. Cells were incubated at 37°C, 5% CO_2_ for a minimum of 2-hours to allow macrophages to adhere. Wells were washed twice with sterile PBS to remove non-adherent cells and new, pre-warmed growth media added. For cells requiring *ex-vivo* stimulation, 100U of IFNγ (R&D) or vehicle (H _2_O) was added for 18- hours. For co-treatments, a final concentration of 10 µM Nocodazole (Sigma), 40 nM Bafilomycin A1 (Sigma), 100nM Torin1 (Calbiochem) and 100 nM PF360 (PF-06685360; MedChem) was used.

### Nucleofection and plating of peritoneal macrophages

Peritoneal macrophages were harvested from mice as previously described. Once passed through 70uM nylon filter, tubes spun at 90 × g for 10 minutes at 4°C. Cells were resuspended and counted as previously described. Cells were aliquoted into 50 mL falcons with 1 × 10^6^ cells per nucleofection reaction. Cells were spun at 90 × g for 10 minutes at 4°C. Supernatant was carefully aspirated so not to disturb the cell pellet, and cells resuspended in nucleofection buffer (acclimated to room temperature; Lonza; P2 Primary Cell 4D-Nucleofector X Kit L; V4XP-2024) containing 1 µM *Lrrk2* or Scramble ASO per 100 µL, to a final concentration of 1 × 10^6^ cells per 100 µL. 100 µL of cells were transferred to each Nucleocuvette, which was then placed into a 4D-Nucleofector® X Unit (Lonza) and pulsed using the CM 138 pulse code. After nucleofection, 400 µL of growth media (acclimated in incubator 1-hour prior) was added to each nucleovette and cells transferred to plates pre-coated with poly-D-Lysine (Sigma). Cells were left to incubate for 24-hours, after which media was aspirated, cells washed and assays started as previously described.

### Flow cytometry

1-hour prior to collection, BMV109 Pan Cathepsin probe (Vergent Bioscience) and DQ red BSA (Invitrogen) were added to each well at a final concentration of 1µM and 10µG/mL, respectively, and cells incubated at 37°C for 1-hour. Cells were then washed 3 times in sterile PBS, harvested, and transferred to a v-bottom 96-well plate (Sigma, CLS3896-48EA) and centrifuged at 300 × *g* for 5 minutes at 4°C. Cells were resuspended in 50 µL of PBS containing diluted fluorophore-conjugated antibodies (see Table 1) and incubated in the dark at 4°C for 20 minutes. Cells were centrifuged at 300 × g for 5 minutes at 4°C and washed in PBS × 2. Cells were fixed in 50 µL of 1% paraformaldehyde (PFA) at 4°C in the dark for 30 minutes. Cells were centrifuged at 300 × *g* for 5 minutes and resuspended in 200 µL FACs buffer (PBS, 0.5 mM EDTA, 0.1% sodium azide). Cells were taken for flow cytometry on a Macs Quant Analyzer (Miltenyi) or BD LSRFortessa™ Cell Analyzer. A minimum of 100,000 events were captured per sample and data were analyzed using FlowJo version 10.6.2 software (BD Biosciences). When validating flow cytometry panels and antibodies, fluorescence minus one controls (FMOCs) were used to set gates and isotype controls were used to ensure antibody-specific binding.

**Table 1.** Flow cytometry monocyte marker antibody panel.

| Target | Conjugate | Antibody<br>Cat# | Dilution | Company |
| --- | --- | --- | --- | --- |
| **CD11b** | PE-Cy7 | 101216 | 1:100 | Biolegend |
| **MHC-II** | APC-Cy7 | 107628 | 1:100 | Biolegend |
| **Live/dead**<br>**stain** | Amcyan | 130113144 | 1:2000 | Fisher |
| **FcR** | - | NC0093774 | 1:100 | Fisher |

### Ea _*(52–68)*_ uptake assay

MHC II Ea chain (Ea) (52–68) peptide (Anaspec) was reconstituted in sterile distilled H _2_O to a final concentrate of 1mG/mL. Once peritoneal macrophages had adhered to plates, 5µG per well was added in growth media. Cells were incubated for 18-hours and taken forward for flow cytometry.

### Cytokine release measurements via Mesoscale discovery electrochemiluminescence

V-PLEX mouse pro-inflammatory panel 1 kit (MSD; K15048D) was used to quantify cytokines in conditioned media from pMacs. Media was diluted 1:1 with MSD kit diluent and incubated at room temperature in the provided MSD plate with capture antibodies for 2 hours as per manufacturer’s instructions. Plates were then washed × 3 with PBS with 0.1% Tween-20 and detection antibodies conjugated with electrochemiluminescent labels were added and incubated at room temperature for another 2 hours. After 3 × washes with PBS containing 0.05% Tween-20, MSD read buffer was diluted to 2x and added, and the plates were loaded into the QuickPlex MSD instrument for quantification. Results were normalized to total live cell counts as measured via flow cytometry.

### Immunoblotting

Media was aspirated and cells washed in PBS and lysed in RIPA buffer (50 mM Tris pH 8, 150 mM NaCl, 1% NP-40, 0.5% NaDeoxycholate, 0.1% SDS). Cell lysates were then centrifuged at 10,000 × *g* for 10 mins at 4 °C. 6X Laemmli sample buffer added (12% SDS, 30% β-mercaptoethanol, 60% Glycerol, 0.012% Bromophenol blue and 375 mM Tris pH 6.8) and samples were reduced and denatured at 95°C for 5 minutes. Samples were loaded into 4- 20% Criterion Tris-HCl polyacrylamide gels (BioRad) alongside Precision plus protein dual-color ladder (Biorad) to determine target protein molecular weight. Electrophoresis was performed at 100 V for ∼60 minutes and proteins transferred to a polyvinylidene difluoride (PVDF) membrane using a Trans-Blot Turbo Transfer System (BioRad) which utilizes Trans-Blot Turbo Midi PVDF transfer packs (BioRad) in accordance to manufacturer’s instructions. Prior to blocking, total protein was measured using Revert total protein stain (Licor) and imaged on the Odyssey FC imaging system (Licor). Membranes were then blocked in 5% non-fat milk in TBS/0.1% Tween-20 (TBS-T) for 1 hour at room temperate and subsequently incubated with primary antibody (see Table 2) in blocking solution overnight at 4°C. Membranes were washed with TBS-T (3 × 5 minutes) and incubated in horseradish peroxidase (HRP)-conjugated secondary antibody (1:1000) (BioRad) in blocking solution for 1 hour. Membranes were washed in TBS-T (3x 5 minutes) and developed using Super signal west femto/pico (Thermo). Membranes were imaged using the Odyssey FC imaging system and quantified using Image Studio Lite Version 5.2 (Licor).

**Table 2.** Antibodies for immunoblotting.

| Target | Antibody<br>Cat# | Dilution | Company |
| --- | --- | --- | --- |
| **LRRK2** | ab133474 | 1:1000 | Abcam |
| **LRRK2 pS935** | ab133450 | 1:1000 | Abcam |
| **LC3** | Z275 | 1:3000 | Cell signalling |
| **RILPL1** | ab302492 | 1:1000 | Abcam |
| **mTOR** | ab134903 | 1:1000 | Abcam |
| **S6k pThr389** | 9205 | 1:1000 | Cell signalling |
| **B-actin** | AM4302 | 1:5000 | Thermo |
| **Revert Total Protein** | 926-11011 | - | Licor |

### Nanostring based mRNA expression analysis of lysosomal and immune-related genes

RNA from approximately 2-4 × 10^6^ cells was isolated. RNase Easy mini kit (Qiagen) was used according to manufacturer’s instructions. Briefly, 10 µL of β-mercaptoethanol was added to every 1 mL of RLT buffer. 350 µL was added to each well and cells homogenized manually with a mini cell scraper. Cell lysate was transferred to an RNAase free Eppendorf and an equal volume of 70% ethanol was added to each sample. Samples were loaded into supplier columns and centrifuged at 10,000 rpm for 30 seconds and flow through discarded. 350 µl RW1 buffer and centrifuged at 10,000 rpm for 15 seconds and flow through discarded. RW1 buffer was added and columns centrifuged at 10000 rpm for 15 seconds. RPE buffer was added to columns, centrifuged for 10,000 rpm for 30 seconds, repeated, and 30 µl RNase free water was added and RNA eluted. RNA concentration was quantified and 260/230 and 260/280 recorded using a spectrophotometer. RIN values were assessed to ensure RNA integrity using Agilent RNA 6000 Nano Kit (Agilent) and Agilent 2100 Bioanalyzer.

Approximately 100□ng of total RNA were hybridized to a Custom Panel for profiling 250 mouse genes within lysosomal, autophagy and inflammatory pathways (Sup.file.1) in a final volume of 15□μl at 65□°C for 22□h according to manufacturer’s protocol (NanoString Technologies, Inc., Seattle, WA, USA). Gene expression profiling was measured on the NanoString nCounter™ system. Hybridized samples were processed on the NanoString nCounter™ Preparation Station using the high-sensitivity protocol, in which excess Capture and Reporter Probes were removed and probe□transcript complexes were immobilized on a streptavidin-coated cartridge and data collected on an nCounter digital analyzer (NanoString), following manufacturer’s instructions.

Background level was determined by mean counts of eight negative control probes plus two standard deviations. Samples that contain <50% of probes above background, or that have imaging or positive control linearity flags, were excluded from further analysis. Probes that have raw counts below background in all samples were excluded from differential expression analysis to avoid false positive results. Data were normalized by geometric mean of housekeeping genes. All statistical analyses were performed on log2 transformed normalized counts.

Pre□processing and normalization of the raw counts was performed with nSolver Analysis Software v4.0 (www.nanostring.com). The 6 spiked□in RNA Positive Control and the 8 Negative controls present in the panel were used to confirm the quality of the run. Data were analysed either by ROSALIND® (https://rosalind.onramp.bio/), with a HyperScale architecture developed by ROSALIND, Inc. (San Diego, CA), or in the nSolver Analysis Software v4.0. Fold changes and pValues are calculated using the optimal method as described in the nCounter® Advanced Analysis 2.0 User Manual. P-value adjustment is performed using the Benjamini-Hochberg method of estimating false discovery rates (FDR). Heatmaps of differentially expressed genes were done using nSolver Analysis Software v4.0.

### Dextran-AF488-labelling of lysosomal tubules and microscopy

pMacs were pulsed with 0.5 mg/mL dextran AlexaFluoro-488 or 564 (Invitrogen) for 1 hour, followed by a pulse period of 2 hours to ensure loading of dextran into lysosomes. Cells were then chased in growth media containing 100U IFN γ or vehicle for 2 hours to induce lysosomal tubule formation. Cells were imaged live using either a Leica THUNDER imager or Zeiss Confocal LSM800 (ICBR, University of Florida) at 60/63 × magnification. Image analysis was performed using Cellprofiler 4.2.5.

### Intracellular and extracellular MHC-II immunostaining and microscopy

Methods for intracellular and extracellular MHC-II immunostaining were adapted from [31]. Cells were washed 3x with DPBS^+/+^ and incubated with 25 ug/mL of APC-MHC-II (Biolegend) in DPBS^+/+^ containing FcR blocking reagent for 30 minutes at room temperature. Cells were then washed 3x with DPBS^+/+^ and fixed by incubation in 4% PFA for 10 minutes at room temperature and then washed 3x with DPBS^+/+^. Fixed cells were permeabilized with permeabilization buffer (eBiosciences) on ice for 15 minutes. 25 ug/mL of PE-610-MHC-II (Biolegend) was spiked in and cells incubated for 30 minutes at room temperature. Cells were then washed 3x in DPBS^+/+^ and incubated in 1 μg/ml DAPI (Life Technologies) for 10 minutes at room temperature in DPBS^+/+^. Cells were imaged using a Leica THUNDER imager at 60 × magnification. Image analysis was performed using Cellprofiler 4.2.5.

### Dextran and Transferrin labelling

pMacs were pulsed with 0.5 mg/mL dextran or transferrin AlexaFluoro-488 (Invitrogen) for 1 hour at 37 °C. Cells were washed 3x with DPBS^+/+^, fixed in 4% PFA for 10 minutes at room temperature and incubated in 1 μg/ml DAPI (Life Technologies) for 10 minutes at room temperature in DPBS^+/+^. Cells were imaged on an EVOS M7000 at 20x magnification. Image analysis was performed using Cellprofiler 4.2.5.

### Statistics and data analysis

Data and statistical analyses were performed using IBM SPSS statistics 27 or GraphPad Prism 9. For assessing differences between groups, data were analyzed by either 1-way or 2- way analysis of variance (ANOVA), or by t-test. In instances when data did not fit parametric assumptions, Kruskal-Wallis non-parametric ANOVA was used. Post-hoc tests following ANOVAs were conducted using Tukey HSD or Bonferroni correction. Two-tailed levels of significance were used and p□<□0.05 was considered statistically significant. Graphs are depicted by means +/- standard error of the mean (SEM).

## Acknowledgements

We thank members of the Tansey lab for useful discussions and edits of the manuscript. We thank Biogen/Ionis for supplying all Lrrk2 and scramble ASOs. We thank the UF Interdisciplinary Center for Biotechnology Research (UF | ICBR) for use of flow cytometry facilities and advice.

## Authors contributions

RLW was responsible for experimental design, performing experiments, plotting and analysing data, data interpretation and drafting and editing manuscript. JRM, HAS, and DAG were responsible for optimizing Lrrk2 ASOs. HAS was responsible for running all MSD-experiments. All authors participated in editing the manuscript.

## Authors’ information

Please contact corresponding author at

---

**Supplementary Figure 1.**
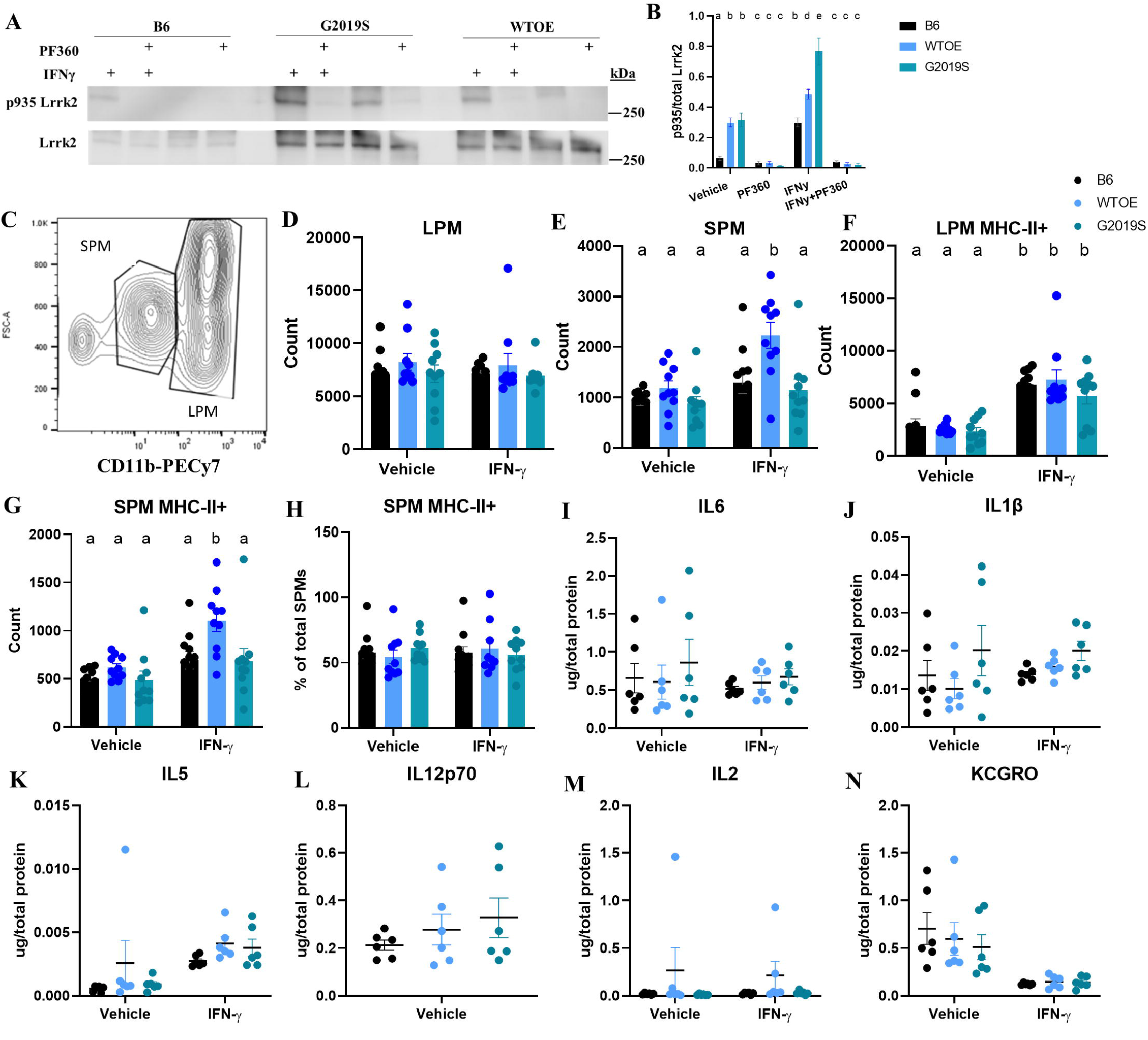
Altered antigen presentation and lysosomal function in G2019S BAC transgenic pMacs: pMacs from 10-12-week-old male B6, WTOE or *G2019S* mice were stimulated with 100U IFNγ +/- 100nM PF360 for 18-hours. (**A, B**) Total Lrrk2 and phosphorylated LRRK2 at S935 were quantified via western blot. Representative western blots shown. (**C**) Cd11b MFI was used to differentiate LPMs from SPMs via flow-cytometry. (**D, E**) LPM and SPM count was quantified. (**F, G**) MHC-II+ LPM and SPM counts were quantified. (**H**) SPM MHC-II+ count was expressed as a % of total SPMs and quantified. (**I, J, K, L, M, N**) Levels of the cytokines IL6, IL1β, IL5, IL12p70, IL2 andKC/GRO in media were assessed, normalized to total protein levels and quantified. Bars represent mean +/- SEM (N = 8-10). Two-way ANOVA, Bonferroni post-hoc, groups sharing the same letters are not significantly different (p>0.05) whilst groups displaying the same letter are significantly different (p<0.05).

**Supplementary Figure 2.**
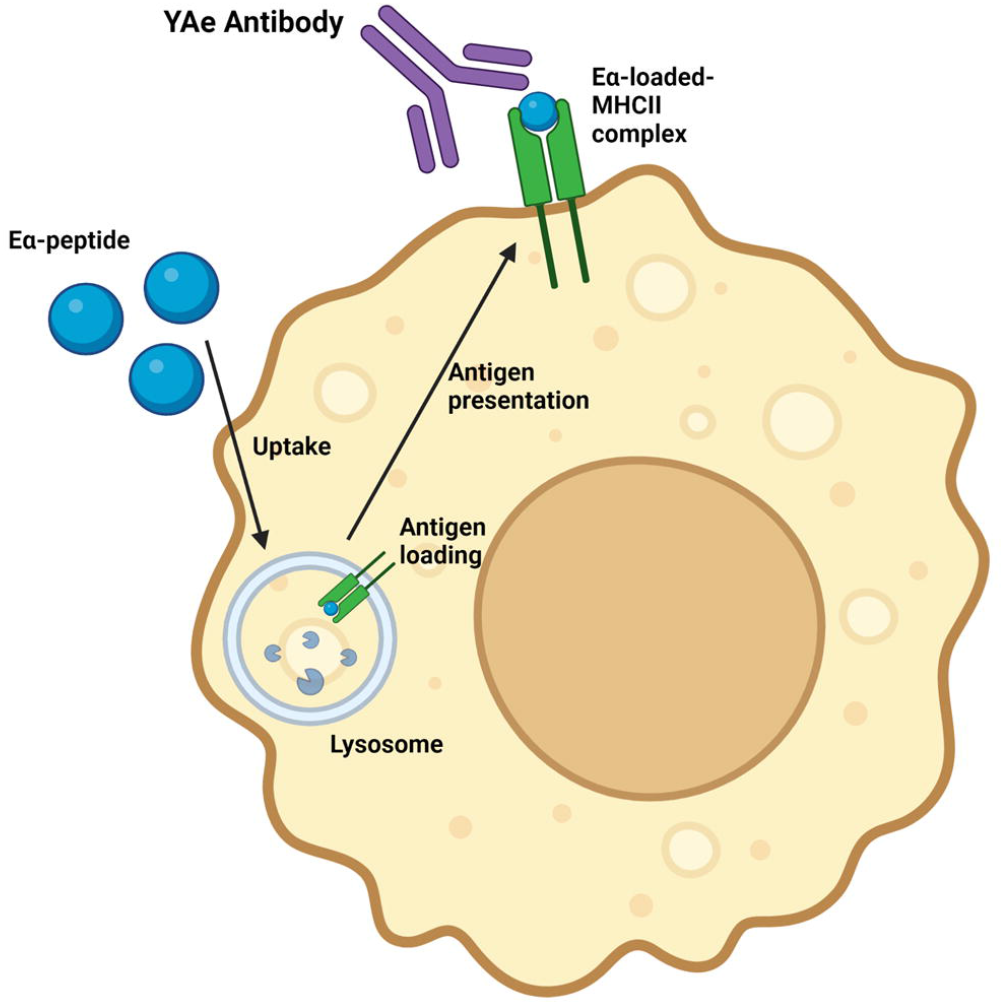
The Eα: YAe model.

**Supplementary Figure 3.**
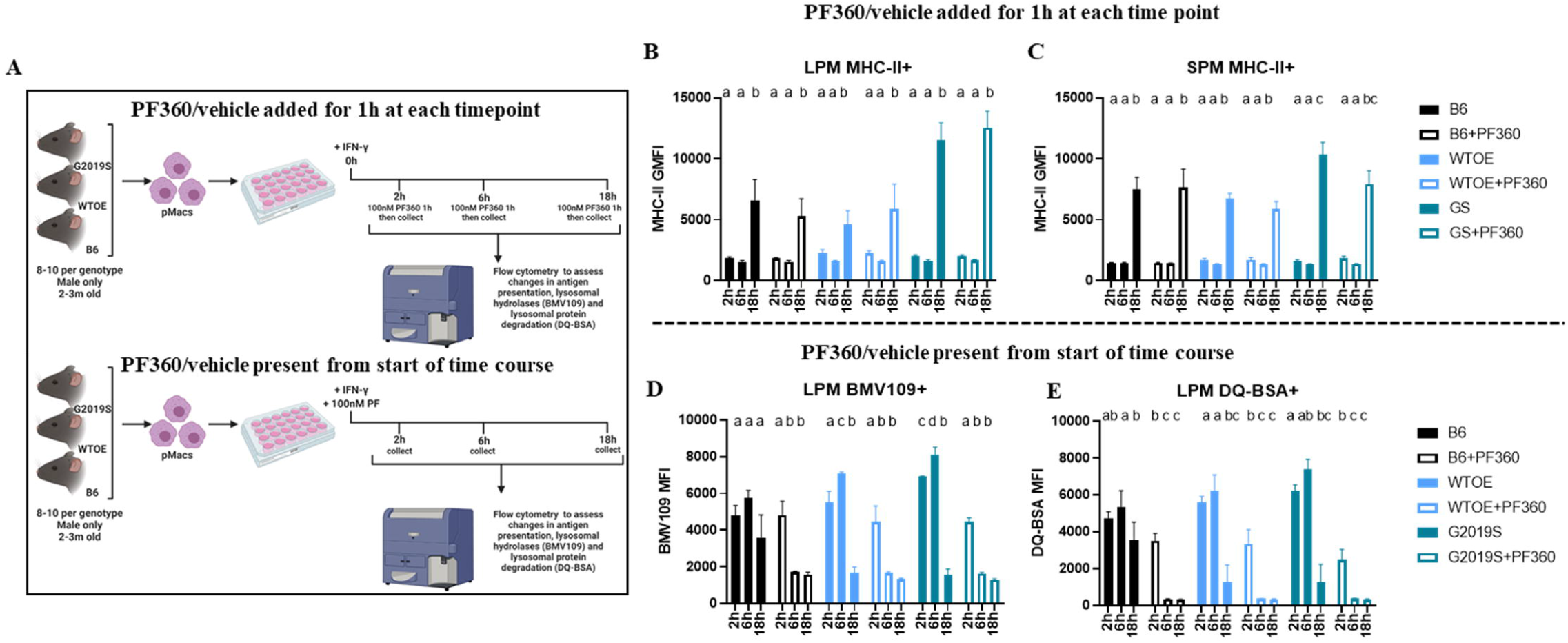
LRRk2 kinase activity modulates lysosomal function early in the inflammatory response in cells engaging in antigen presentation: (A) pMacs from 10-12-week-old male B6, WTOE or G2019S mice were subject to one of two treatment conditions. All cells were stimulated with 100U IFNγ and harvested at 2-, 6- or 18- hours. In the first treatment condition, cells were treated +/- 100nM PF360 from the beginning of the 18-hour treated. In the second treatment condition, cells were treated for 1- hour with 100nM PF360 immediately prior to harvesting. (**B**) LPM MHC-II+ GMFI was quantified via flow cytometry in cells treated with PF360 for 1-hour. (**C**) SPM MHC-II+ GMFI was quantified via flow cytometry DQ-BSA MFI was quantified in LPMs via flow cytometry in cells treated with PF360 for 1-hour. (**D**) BMV109 MFI was quantified in LPMs via flow cytometry in cells treated with PF360 for the duration of the IFNy treatment. (**E**) DQ-Red BSA MFI was quantified in LPMs via flow cytometry in cells treated with PF360 for the duration of the IFNy treatment. Bars represent mean +/- SEM (N = 8-10). Three-way ANOVA, Bonferroni post-hoc, groups sharing the same letters are not significantly different (p>0.05) whilst groups displaying the same letter are significantly different (p<0.05). (**F**) Schematic of hypothesized model: modulation of antigen presentation in pMacs by LRRK2 via regulation of lysosomal activity early in inflammatory response.

**Supplementary Figure 4.**
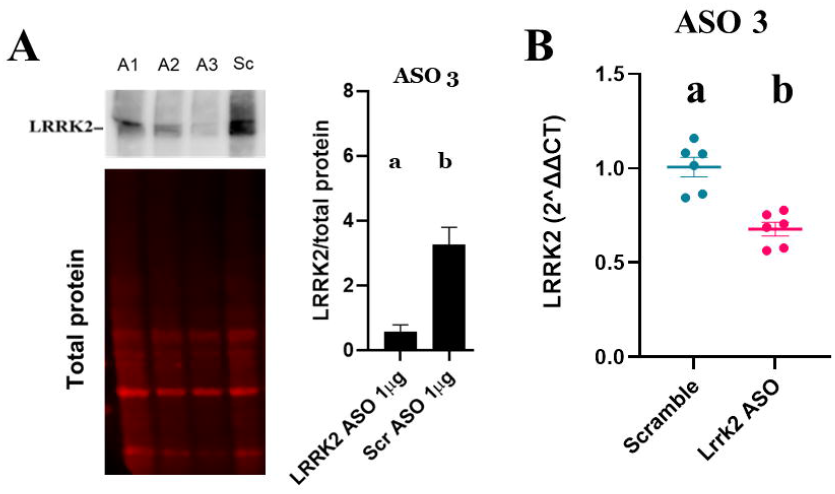
G2019S pMac antigen presentation and lysosomal phenotypes are rescued by knock-down of LRRK2: pMacs from 10-12-week-old male B6, *WTOE* or *G2019S* mice were nuclefoected with 1uG of 1 of 3 Lrrk2-targetting ASOs or a scramble control. (**A**) Total LRRK2 levels were normalized to total protein levels and quantified. Representative western blots shown. (**B**). LRRK2 mRNA levels were quantified, normalized to house-keeping gene expression and expressed as 2^ΔΔCT and fold-change from scramble ASO treated cells. Bars represent mean +/- SEM (N = 5-6). Student’s T-test, groups sharing the same letters are not significantly different (p>0.05) whilst groups displaying the same letter are significantly different (p<0.05).

**Supplementary Figure 5.**
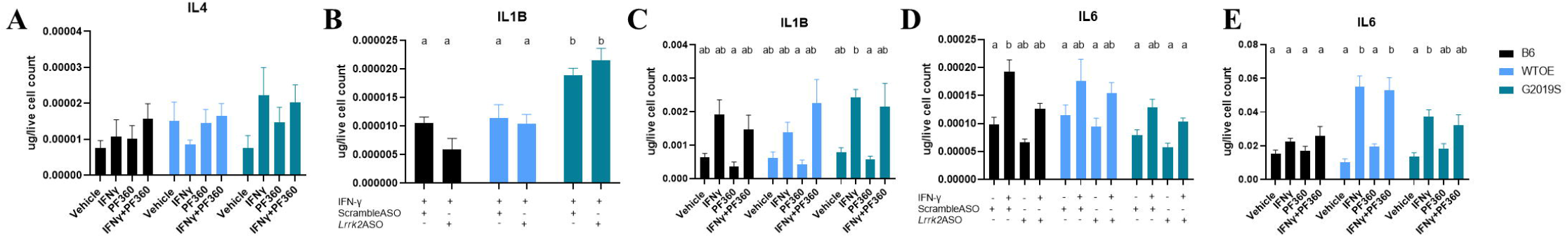
LRRK2 knock-down via antisense oligonucleotide and kinase inhibition alters cytokine release from pMacs: pMacs from 10-12-week-old male B6, WTOE or G2019S mice were nuclefoected with a Lrrk2-targetting ASO or scramble control and stimulated with 100U IFNγ, or were plated with 100U IFNγ +/- 100nM of Pf360 and media collected after 18-hours. Cytokine levels of IL4 (**A**), IL1β (**B,C**), and IL6 (**D, E**) were quantified and normalized to live cell count. Lrrk2 protein levels were assessed and normalized to total protein levels and quantified. Bars represent mean +/- SEM (N = 8-10). Two-way ANOVA, Bonferroni post-hoc, groups sharing the same letters are not significantly different (p>0.05) whilst groups displaying the same letter are significantly different (p<0.05).

**Supplementary figure 6.**
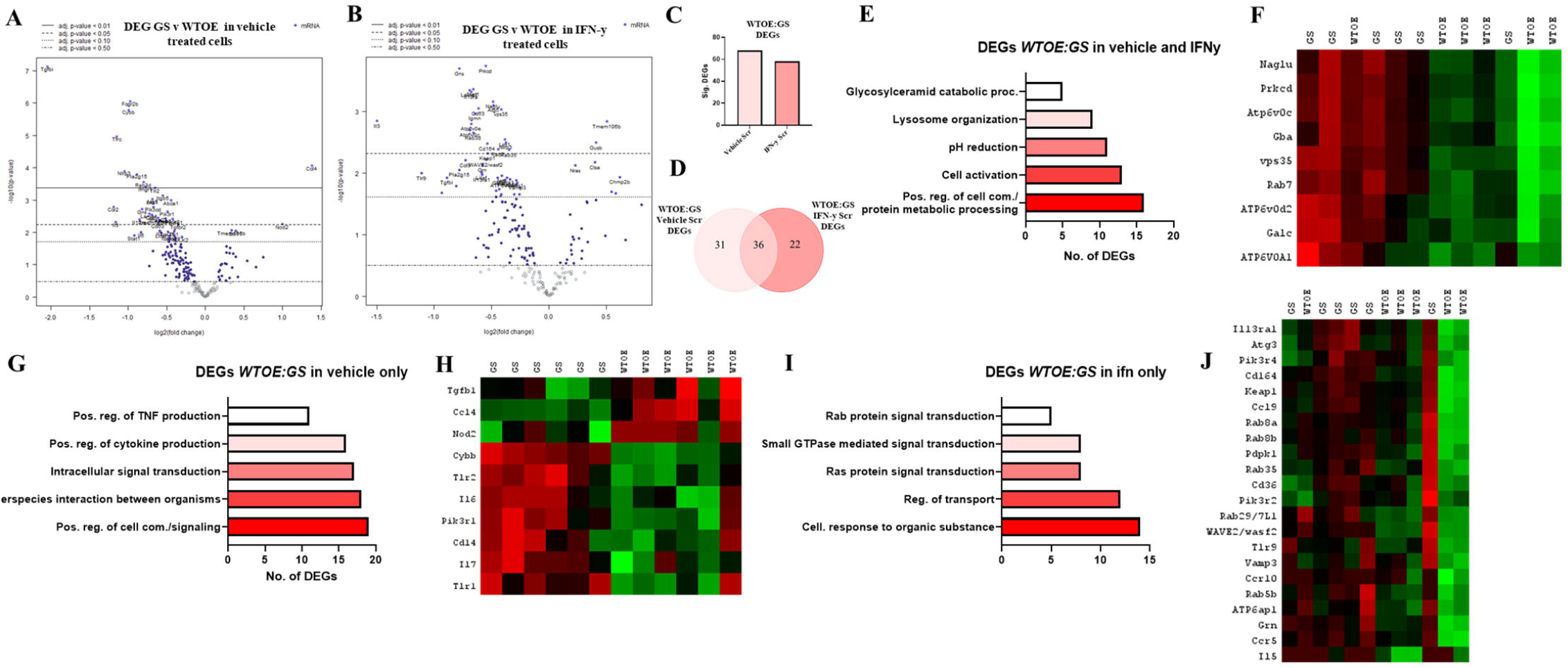
Nanostring-based transcriptome analysis reveals genotype differences in a treatment specific manner and reveals differential response to IFN-γ by G2019S pMacs: Transcriptomic analysis from vehicle (**A**) or IFNγ (**B**) treated *G2019S* and *WTOE* pMacs Volcano plot shows proteins with fold change□>□1.5 and an adjusted p-value□≤□0.05. (**C, D**) Significant DEGs were counted and compared across treatments. (**E**) ShinyGO 0.76.3 was used to identify pathways in which significant DEGs were associated with. (**F**) Heat maps show DEGs in both vehicle and IFNγ treatment. (**G**) ShinyGO 0.76.3 was used to identify pathways in which significant DEGs were associated with. (**H**) Heat maps show DEGs in only vehicle treatment. (**I**) ShinyGO 0.76.3 was used to identify pathways in which significant DEGs were associated with. (**J**) Heat maps show DEGs in only IFNγ treatment.

**Supplementary Figure 7.**
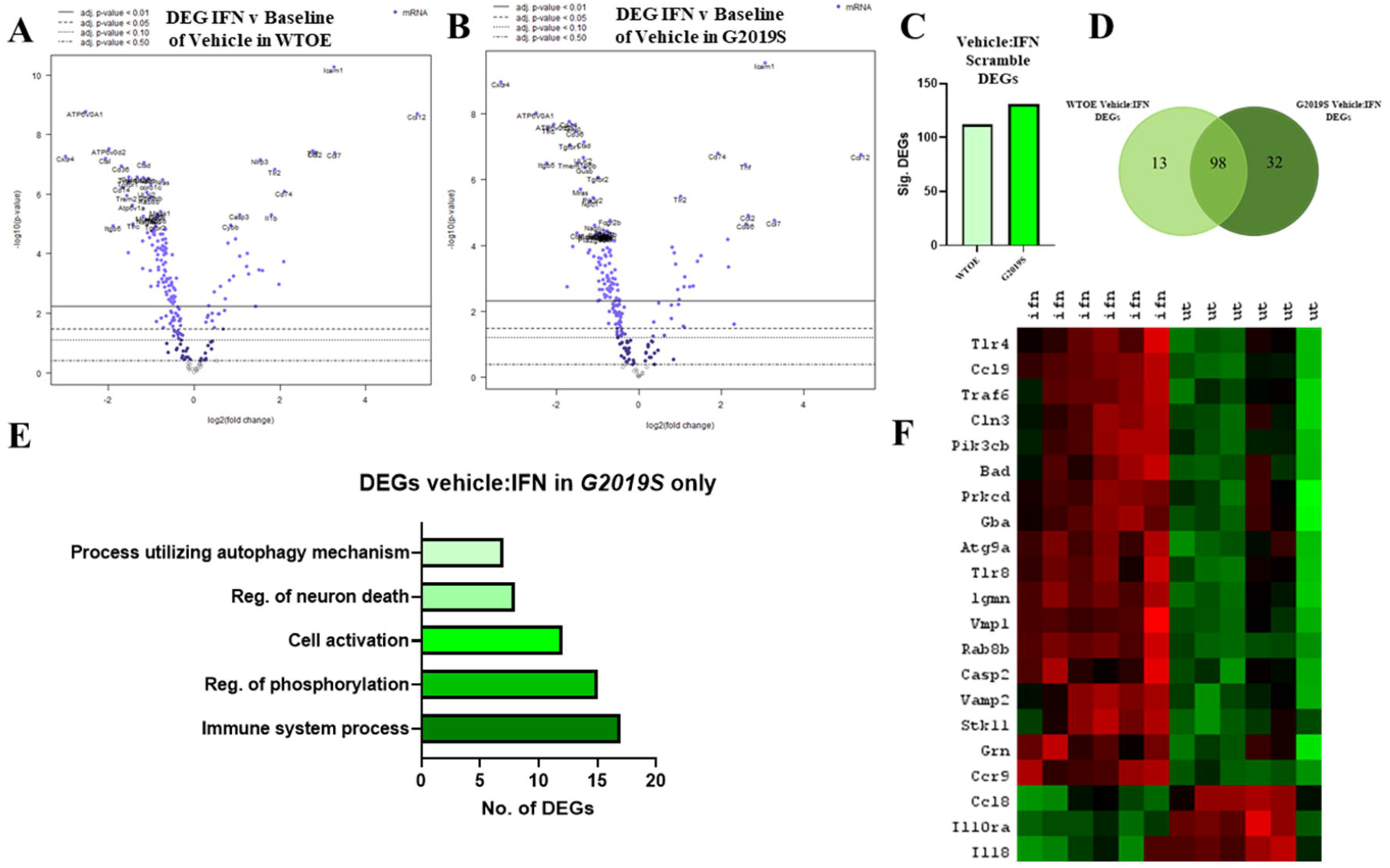
Nanostring-based transcriptome analysis reveals differential response to IFN-γ by G2019S pMacs: Transcriptomic analysis from *WTOE* (**A**) or *G2019S* (**B**) vehicle and IFNγ-treated pMacs. Volcano plot shows proteins with fold change□>□1.5 and an adjusted p-value□≤□0.05. (**C, D**) Significant DEGs were counted and compared across genotypes. (**E**) ShinyGO 0.76.3 was used to identify pathways in which significant DEGs were associated with. (**F**) Heat maps show DEGs seen only in *G2019S* pMacs.

**Supplementary Figure 8.**
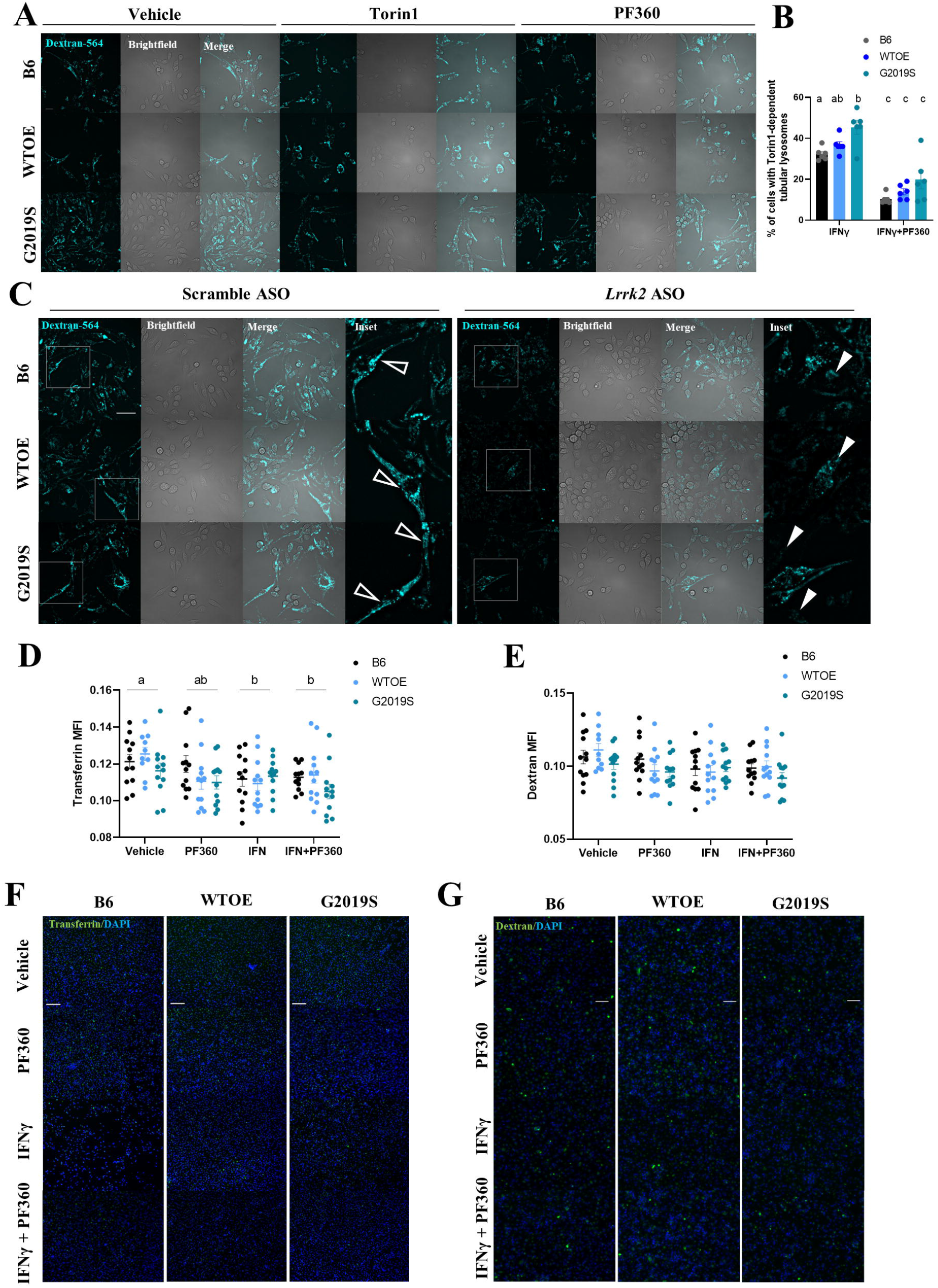
LRRK2 modulates antigen presentation via lysosomal tubule formation: (A) pMacs from 10-12-week-old male B6, WTOE or *G2019S* mice treated with 0.5mg/mL Dextran Alexa-Fluor546 for 1-hour, followed by a 2-hour pulse-period to ensure loading into lysosomes, treated with 100U of IFNy +/- 100nM Torin1 or 100nM PF360 for 2-hours to stimulate LTF and imaged live. (**B**) Percentage of cells with Torin1-dependent tubular lysosomes was quantified. (**C**) pMacs from 10-12-week-old male B6, WTOE or *G2019S* mice were nucleofected with 1uG of scramble or Lrrk2-targetting ASO, allowed to rest 24 hours, then treated with 0.5mg/mL Dextran Alexa-Fluor546 for 1-hour, followed by a 2-hour pulse-period to ensure loading into lysosomes, treated with 100U of IFNy to stimulate LTF and imaged live. Filled white arrows indicate pMacs with tubular structures, empty arrow heads indicate pMacs with punctate dextran. Scale bars, 10μM (**D, E, F, G**) pMacs were loaded with 0.5mG/mL of Dextran or transferrin Alexa-Fluor488 for 1 hour, fixed, imaged and uptake quantified. Bars represent mean +/- SEM (N = 4-6). Two-way ANOVA, Bonferroni post-hoc, groups sharing the same letters are not significantly different (p>0.05) whilst groups displaying the same letter are significantly different (p<0.05). Scale bars, 40μM.

**Supplementary Figure 9.**
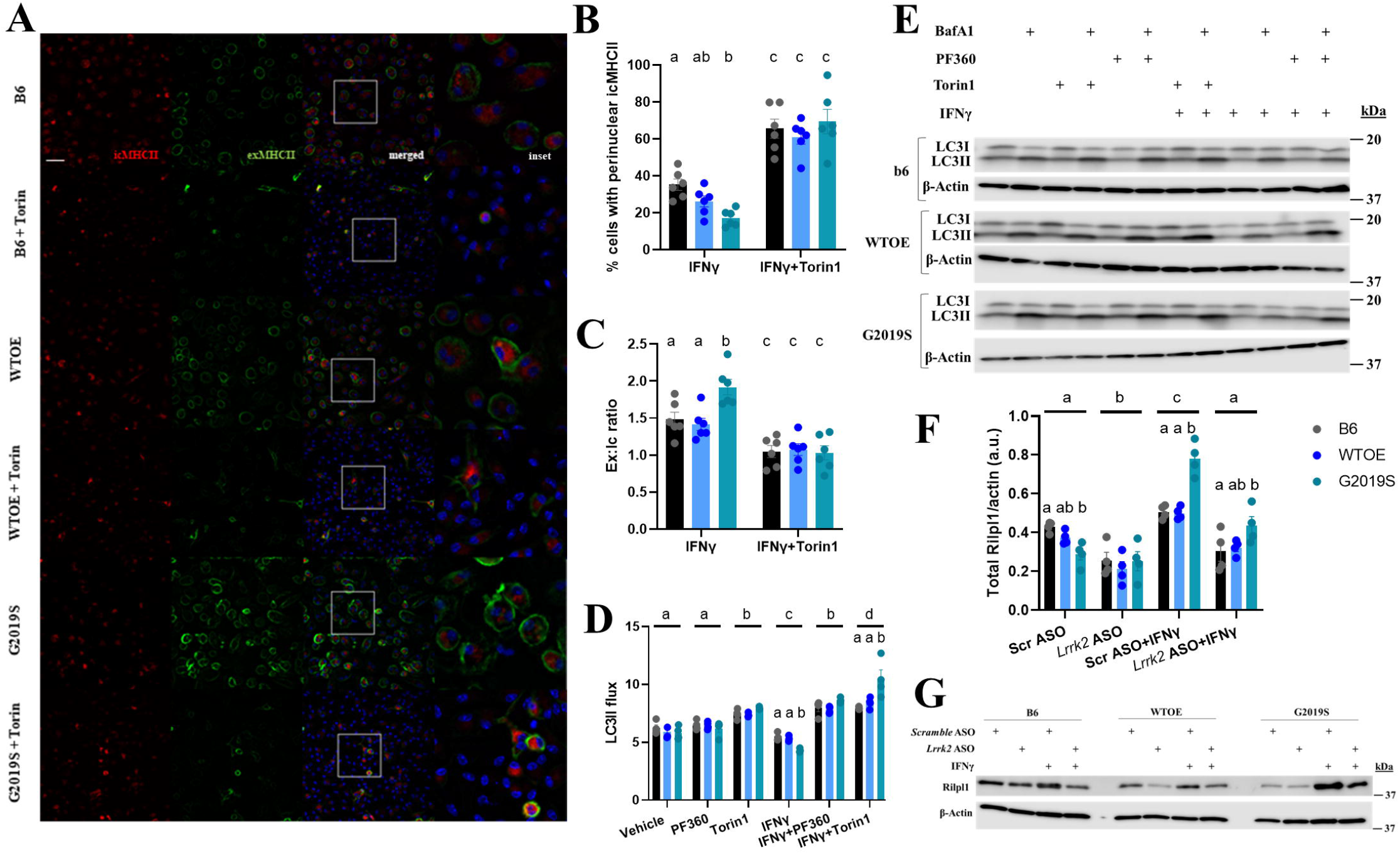
LRRK2 modulates MHC-II trafficking and autophagic flux : pMacs from 10-12-week-old male B6, WTOE or *G2019S* mice were treated with 100U of IFNy +/- 100nM Torin1 for 18-hours and stained for intracellular and extracellular MHC-II and Ex:Ic ratio quantified and perinuclear clustering % quantified. Scale bars, 30μM. (**A, B, C**). pMacs were treated with 100U IFNy +/- 100nM Torin1 or PF360 for 18-hours, with 40nM Bafilomycin A1 added to final 2-hours of treatment. Protein lysate quantified for LC3-II levels and LC3 flux quantified (**D, E**). Representative western blots shown. pMacs were treated with 100U of IFNy +/- 100nM PF360 and protein lysate assessed for RILPL1 protein levels and normalized to β-actin levels and quantified. Representative western blots shown. Bars represent mean +/- SEM (N = 4-6). Two-way ANOVA, Bonferroni post-hoc, groups sharing the same letters are not significantly different (p>0.05) whilst groups displaying the same letter are significantly different (p<0.05).

